# Structural phylogenetics unravels the evolutionary diversification of communication systems in gram-positive bacteria and their viruses

**DOI:** 10.1101/2023.09.19.558401

**Authors:** David Moi, Charles Bernard, Martin Steinegger, Yannis Nevers, Mauricio Langleib, Christophe Dessimoz

**Affiliations:** Department of Computational Biology, University of Lausanne, Lausanne, Switzerland; Swiss Institute of Bioinformatics, Lausanne, Switzerland; School of Biological Sciences, Seoul National University, Seoul, South Korea; Artificial Intelligence Institute, Seoul National University, Seoul, South Korea; Institute of Molecular Biology and Genetics, Seoul National University, Seoul, South Korea; Unidad de Bioinformática, Institut Pasteur de Montevideo, Montevideo, Uruguay; Unidad de Genómica Evolutiva, Facultad de Ciencias, Universidad de la República, Montevideo, Uruguay

## Abstract

Recent advances in AI-based protein structure modeling have yielded remarkable progress in predicting protein structures. Since structures are constrained by their biological function, their geometry tends to evolve more slowly than the underlying amino acids sequences. This feature of structures could in principle be used to reconstruct phylogenetic trees over longer evolutionary timescales than sequence-based approaches, but until now a reliable structure-based tree building method has been elusive. Here, we introduce a rigorous framework for empirical tree accuracy evaluation and tested multiple approaches using sequence and structure information. The best results were obtained by inferring trees from sequences aligned using a local structural alphabet—an approach robust to conformational changes that confound traditional structural distance measures. We illustrate the power of structure-informed phylogenetics by deciphering the evolutionary diversification of a particularly challenging family: the fast-evolving RRNPPA quorum sensing receptors. We were able to propose a more parsimonious evolutionary history for this critical protein family which enables gram-positive bacteria, plasmids and bacteriophages to communicate and coordinate key behaviors. The advent of high-accuracy structural phylogenetics enables a myriad of applications across biology, such as uncovering deeper evolutionary relationships, elucidating unknown protein functions, or refining the design of bioengineered molecules.

## Introduction

Since Darwin, phylogenetic trees have depicted evolutionary relationships among organisms, viruses, genes, and other evolving entities, allowing us to understand their shared ancestry and tracing the events that led to the observable extant diversity. Trees based on molecular data are typically reconstructed from nucleotide or amino-acid sequences, by aligning homologous sequences and inferring the tree topology and branch lengths under a model of character substitution^1–3^. However, over long evolutionary time scales, multiple substitutions occurring at the same site cause uncertainty in alignment and tree building. The problem is particularly acute when dealing with fast evolving sequences, such as viral or immune-related ones, or when attempting to resolve distant relationships, such as at the origins of animals^4–6^ or beyond.

In contrast, the fold of proteins is often conserved well past sequence signal saturation. Furthermore, because 3D structure determines function, protein structures have long been studied to gain insight into their biological role within the cell, whether it be catalyzing reactions, interacting with other proteins to form complexes or regulating the expression of genes, among others. Until recently, protein structures had to be obtained through labor-intensive crystallography and other experimental methods, with modeling efforts often falling short of the level of accuracy required for the many tasks structures were used for. Due to these limitations, structural biology and phylogenetics have developed as largely separate disciplines, and each field has created different models describing evolutionary or molecular phenomena suited to the availability of computational power and experimental data. Despite these limitations, attempts have been made to merge the two paradigms ^7–9^.

Now, the widespread availability of accurate structural models based on AI predictions^10,11^ opens up the prospect of reconstructing trees from structures. However, there are pitfalls to avoid in order to derive evolutionary distances between homologous protein structures. Geometric distances between rigid body representations of structures, such as root mean square deviation (RMSD) distance or template modeling (TM) score^12^, are confounded by spatial variations caused by conformational changes^13,14^. More local structural similarity measures have been proposed in the context of protein classification^13^, but due to the relative paucity of available structures until recently, little is known about the accuracy of structure-based phylogenetic reconstruction beyond a few isolated case studies^15,16^.

Here, we report a comprehensive evaluation of phylogenetic trees reconstructed from the structures of thousands of protein families across the tree of life, using multiple kinds of distance measures. We tested multiple structure-informed approaches, using divergence measures obtained using Foldseek^17^, which outputs scores from rigid body alignment, local superposition-free alignment and structural alphabet-based sequence alignments. In addition, we tested a recently proposed partitioned structure and sequence likelihood method^18^. The performance of these approaches has been previously assessed on the task of detecting whether folds are homologous and belong to the same family^17,19,20^, or on a few examples^18^, but have never been systematically evaluated for phylogenetic tree inference. Remarkably, we found that some, though not all, structure-informed approaches are competitive with state-of-the-art sequence based phylogenetic methods, and outperform them on highly divergent datasets across benchmarks related to tree topology (TCS and ASTRAL-based species tree branch support) as well as testing the adherence to a molecular clock.

To demonstrate the capabilities of structural phylogenetics, we employ the currently best approach, released as open-source software named *FoldTree*, to resolve the difficult phylogeny of a fast-evolving protein family of high relevance: the RRNPPA (Rap, Rgg, NprR, PlcR, PrgX and AimR) receptors of communication peptides. Although these receptors were identified in the early 1990s^21,22^, their evolutionary history is unclear due to frequent mutations and transfers, making sequence comparisons challenging^23–25^. This is reflected by the nomenclature of the family which were historically described as six different families of intracellular receptors, and of which only structural comparisons allowed to establish the actual consensus on their common evolutionary origin^25–27^. These proteins allow gram-positive bacteria, their plasmids, and their viruses to assess their population density and regulate key biological processes accordingly. These communication systems have been shown to regulate virulence, biofilm formation, sporulation, competence, solventogenesis, antibiotic resistance or antimicrobial production in bacteria^28–32^, conjugation in conjugative elements, lysis/lysogeny decision in bacteriophages^33^ and host manipulation by mobile genetic elements (MGEs)^30,34^. These receptors are paired with a small secreted communication peptide that accumulates extracellularly as the encoding population replicates. Once a quorum of cells, plasmids or viruses is met, communication peptides get frequently internalized within cells and binds to the tetratricopeptide repeats (TPRs) of cognate intracellular receptors, leading to gene or protein activation or inhibition, facilitating a coordinated response beneficial for a dense population. Accordingly, the RRNPPA family impacts human health as it connects to the virulence and transmissibility of pathogenic bacteria and the spread of antimicrobial resistance genes through horizontal gene transfers. We analyze and discuss the relative parsimony of the phylogeny of this family, highlighting the contrasts with the sequence-based tree.

## Results

### Structure-informed trees can outperform sequence-only trees

To incorporate structural information in phylogenetic tree building, we investigated the use of local superposition-free comparison (local distance difference test; LDDT^20^), rigid body alignment (TM score^12^) and a distance derived from a statistically corrected sequence similarity after aligning with a structural alphabet (Fident)^17^. These measures were used to compute distance trees using neighbor joining, after being aligned in an all-vs-all comparison using the Foldseek structural alphabet (*Methods*).

Assessing the accuracy of trees reconstructed from empirical data is notoriously difficult. We used two complementary indicators, ‘correct’ topology and adherence to a molecular clock. We designed the taxonomic congruence score (TCS) (*Methods* and **Supp. Figures 1 and 3**), to assess the congruence of reconstructed protein trees with the known taxonomy^35^. Among several potential tree topologies reconstructed from the same set of input proteins, the ‘better’ topologies can be expected to have higher TCS on average. This is due to the fact that in the majority of cases, gene families are inherited vertically in cellular organisms from parent to daughter cells^36^. The TCS metric used in these benchmarks is designed to weigh topological congruence closer to the root more heavily than towards the leaves.

For trees reconstructed from closely related protein families using standard sequence alignments (the ‘OMA dataset’, see methods), we tested a battery of methods using combinations of structure and sequence input data paired with either multiple sequence alignment (MSA) and maximum likelihood (ML) approaches to build trees or distance based neighbor joining tree building (NJ) (**Figure 1a and methods)**. Trees derived from the combination of sequence and structural alignment, based on the statistically corrected Fident distance (henceforth referred to as the *FoldTree* approach) outperformed other approaches in garnering the highest percentage of top scoring trees within this dataset (**Figure 1b**). This trend was observed across various protein family subsets, taken from taxonomically defined clades within the species tree at varying evolutionary distances (**Supp. Figure 11).** We also experimented with other parameter variations, but they did not lead to further improvements (**Supp. Figures 6,7,8).** In particular, we did not see an improvement when we used only the 3Di structural alphabet in both the alignment and distance estimation steps using both distance and maximum likelihood tree building strategies, indicating that our use of structures mainly contributes to better identifying the homologous residues (**Supp. Figure 6**).

**Figure 1.**
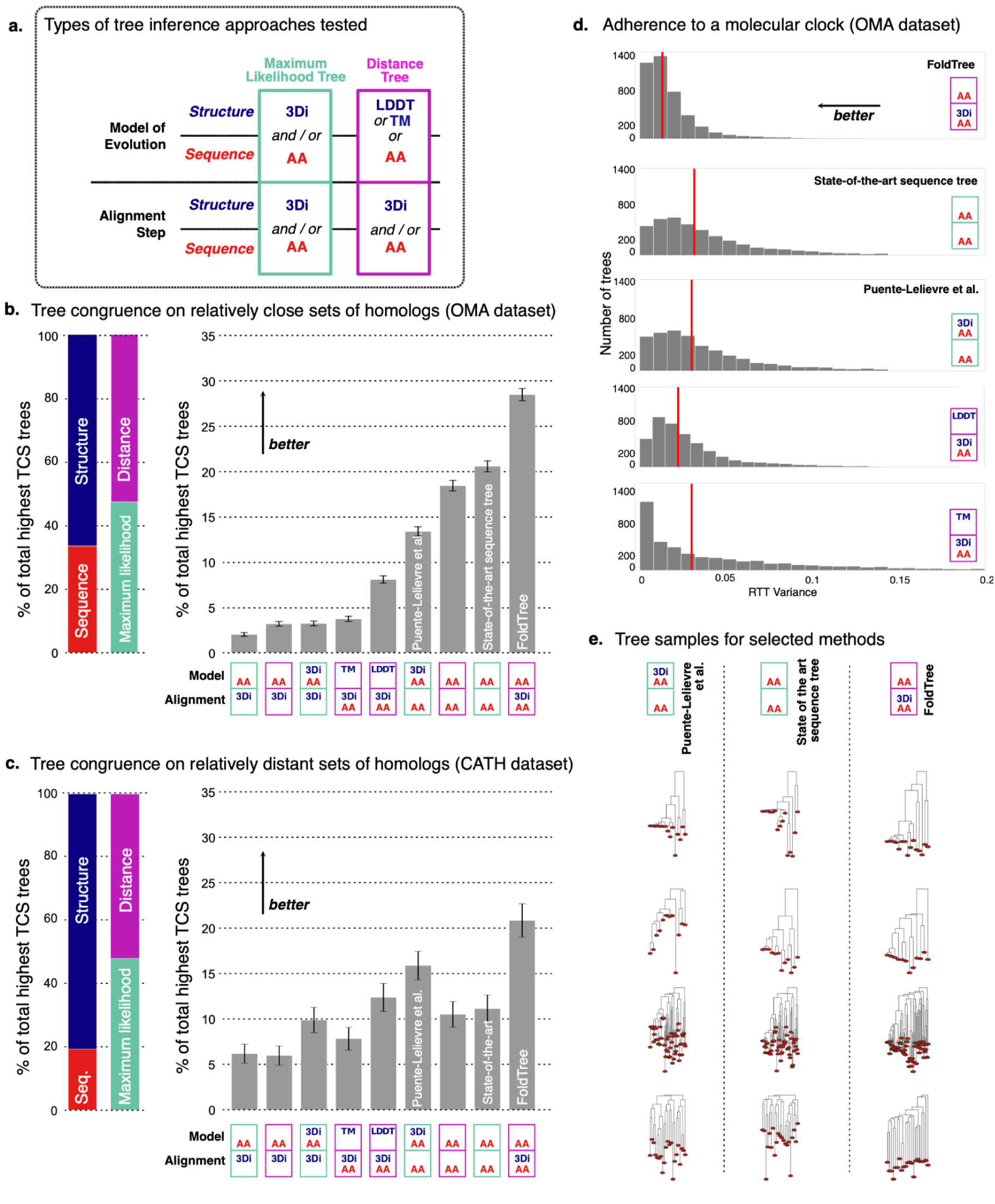
**a)** A diverse set of tree building methods using amino-acid and/or structure information were tested (*Methods* and *Supplementary Information*) **b)** Among the approaches tested, the on incorporating structure information in the alignment phase (“FoldTree”) exhibited the highest proportion of trees with highest Taxonomic Congruence Score (TCS) on the OMA dataset (protein families defined from sequences; relatively close). By contrast, structure trees from LDDT and TM underperform sequence trees and maximum likelihood trees built with partition models of sequence and structural characters. Error bars are derived from a hypergeometric distribution with the same number of samples and classes as the dataset **c)** In the CATH dataset of structurally defined protein families, FoldTree trees metric garner a higher proportion of the trees with the highest TCS per family. The proportion of highest scoring trees using structurally informed methods is greater than in the families defined in the OMA dataset. Error bars are derived from a hypergeometric distribution with the same number of samples and classes as the dataset **d)** The variance of normalized root-to-tip distances were compiled for all trees within the OMA dataset for all tree structural tree methods and sequence trees. FoldTree has a lower mean variance than other methods. The mean of each distribution is shown with a vertical red line. Distributions are truncated to values between 0 and 0.2; **e)** A random sample of trees is shown where each column is from from equivalent protein input sets and each column of trees is derived using a distinct tree building method

We then compared FoldTree with other methods over larger evolutionary distances, using structure-informed homologous families from the CATH database^37^. This database classifies proteins hierarchically, grouping them based on **C**lass, **A**rchitecture, **T**opology and **H**omology of experimentally determined protein structures. We examined sets of proteins from the same homology set after filtering for redundancy (*Methods*). Efforts were made to correct crystal structures with discontinuities or other defects before treebuilding (*Methods*) since these adversely affect structural comparisons. When measuring TCS in families from this more divergent CATH dataset, structure-based methods performed better overall. FoldTree outperformed the sequence-based methods by a larger margin and structurally informed trees received a larger proportion of highest scoring trees overall (**Figure 1c**). In particular, structurally informed maximum likelihood trees^18^ also benefited relative to purely sequence-based methods. Still, in a pairwise comparison between this method and FoldTree, the simpler evolutionary model of FoldTree outperformed this approach in terms of TCS (**Supp. Figure 9**). Additional pairwise comparisons between structural distance methods and maximum likelihood trees show that structural methods provide more congruent topologies with CATH protein families which allow for greater evolutionary distances (**Supp. Figure 13**).

Using both the CATH and OMA datasets, we performed an additional benchmark, recently proposed^38^ as a reaction to an earlier version of this work. Their benchmark is conceptually similar to TCS, but puts more weight on topological differences to the reference species tree close to the leaves (**Supp. Methods)**. This benchmark yielded generally consistent results (**Supp. Figures 14-15)** but on the closer families defined by OMA, FoldTree is slightly outperformed by a sequence-based maximum likelihood approach^38^. However, in more divergent families defined by CATH, FoldTree retains an advantage over that package (**Supp. Figure 15**).

We found that filtering input families based on the confidence of the AlphaFold structure prediction (predicted LDDT or pLDDT) increased the proportion of trees where FoldTree was outperforming maximum likelihood models (**Supp. Figure 12**). This suggests that advancements in structural prediction might further benefit structural trees in the future. We also characterized the distributions of pairwise identities within the induced pairs of sequence-based and 3di-based MSAs as well as Foldseek-based MSAs which combine both alphabets. Here we found that the distribution of mean pairwise percentage identity within alignments was increased by the use of Foldseek and leveraging structure and sequence in tandem to align residues **(Supp. Figure 4-5)**. We also investigated the relationship between sequence divergence within families and the observed differences in TCS between FoldTree and maximum likelihood approaches. We found that as mean amino acid percent identity decreases, the proportion of Foldtree trees with better topologies than the maximum likelihood trees increases **(Supp. Figure 12)**.

To validate our findings using another independent indicator of tree quality, we also assessed how uniform a tree’s root-to-tip lengths are for all its tips, akin to following a molecular clock. Although strict adherence to a molecular clock is unlikely in general, it is reasonable to assume that distance measures resulting in more monotonic trees on average (i.e., with reduced root-to-tip variance, see *Methods*) are more accurate^39^. However, this in and of itself is not a strong indicator of tree quality and must be considered in conjunction with correct topology using a metric such as TCS or ASTRAL quarter support values. We found that in the OMA sequence-based family dataset, FoldTree had the lowest root-to-tip variance of all approaches (**Figure 1d**). The difference is observable in visual comparison of tree shapes for several randomly chosen families (**Figure 1e**). The uniformity of FoldTree root to tip lengths appears despite the higher variance in percent identity seen across the distribution of pairwise alignment in all families (**Supp. Figures 4, 5, 8, 13**). Again, when comparing to maximum likelihood trees informed by structural characters^18^, we see that adding structural information also helps creating trees with more regular root to tip lengths but branch lengths still have more variance than the statistically corrected pairwise distances used by FoldTree.

Both orthogonal metrics of adherence to a molecular clock and species tree discordance indicate that FoldTree produces trees with desirable characteristics that are ideal for constructing phylogenies with sets of highly divergent homologs. However, when considering finer grained topological differences to the species taxonomy with ASTRAL, we do observe a slightly worse performance in the OMA dataset and an equivalent performance in the CATH dataset (**Supp. Figures 14-15**). These results imply that FoldTree is best suited to resolving deeper evolutionary relationships and may suffer in quality compared to maximum likelihood approaches closer to the leaves. We stress that assessing tree quality empirically without relying on an underlying model is difficult and individual phylogenies generated by any model should be interpreted using independent sources of information in order to verify the relative parsimony of each topology between tree inference approaches in and the implied evolutionary scenario recapitulating the extant observable biology. Here we present one such case study centered on the RRNPPA peptidic quorum sensing systems.

### FoldTree reveals the evolutionary diversification of RRNPPA communication systems

To illustrate the potential of structural phylogenies, we reconstructed the evolutionary history of the RRNPPA family of intracellular quorum sensing receptors in gram-positive Bacillota bacteria, their conjugative elements and temperate bacteriophages^25,28,32^. Recently, pioneering work combining structural comparisons among folds and sequence-based phylogenetics have provided insights among some of these families^25^, but a comprehensive reconstruction of the evolutionary history of this family that includes all described subfamilies^30^ remains elusive.

RRNPPA receptors have a core architecture composed of 5 TPRs, and may optionally harbor a HTH DNA-binding domain at the N-terminus and 4 additional TPRs in the C-terminus ^25^. To avoid domain architecture changes confounding the phylogenetic signal, we applied the corecut preprocessing pipeline detailed in methods and **Supp. Figure 2**. We then derived a FoldTree phylogeny for all of the shared core domains of the RRNPPA family. The unrooted tree delineates three main clans. The clan A is composed of only 5 TPRs folds (PlcR, TprA, PrgX, TraA, ComR and Rgg validated subfamilies). The clan B encompasses the non-degenerated 9 TPRs folds (QssR, NprR and Rap validated subfamilies). Finally, the clan C encompasses 9 TPRs folds harboring more degenerated TPR motifs (AloR and AimR validated subfamilies) (**Figure 2a and 2d**). This is more parsimonious than the scenario implied by the sequence-based tree, in which clans of 5 TPRs folds and 9 TPRs are interleaved and suggest a less plausible convergent evolution of protein architectures (**Supp. Figure 16**).

**Figure 2.**
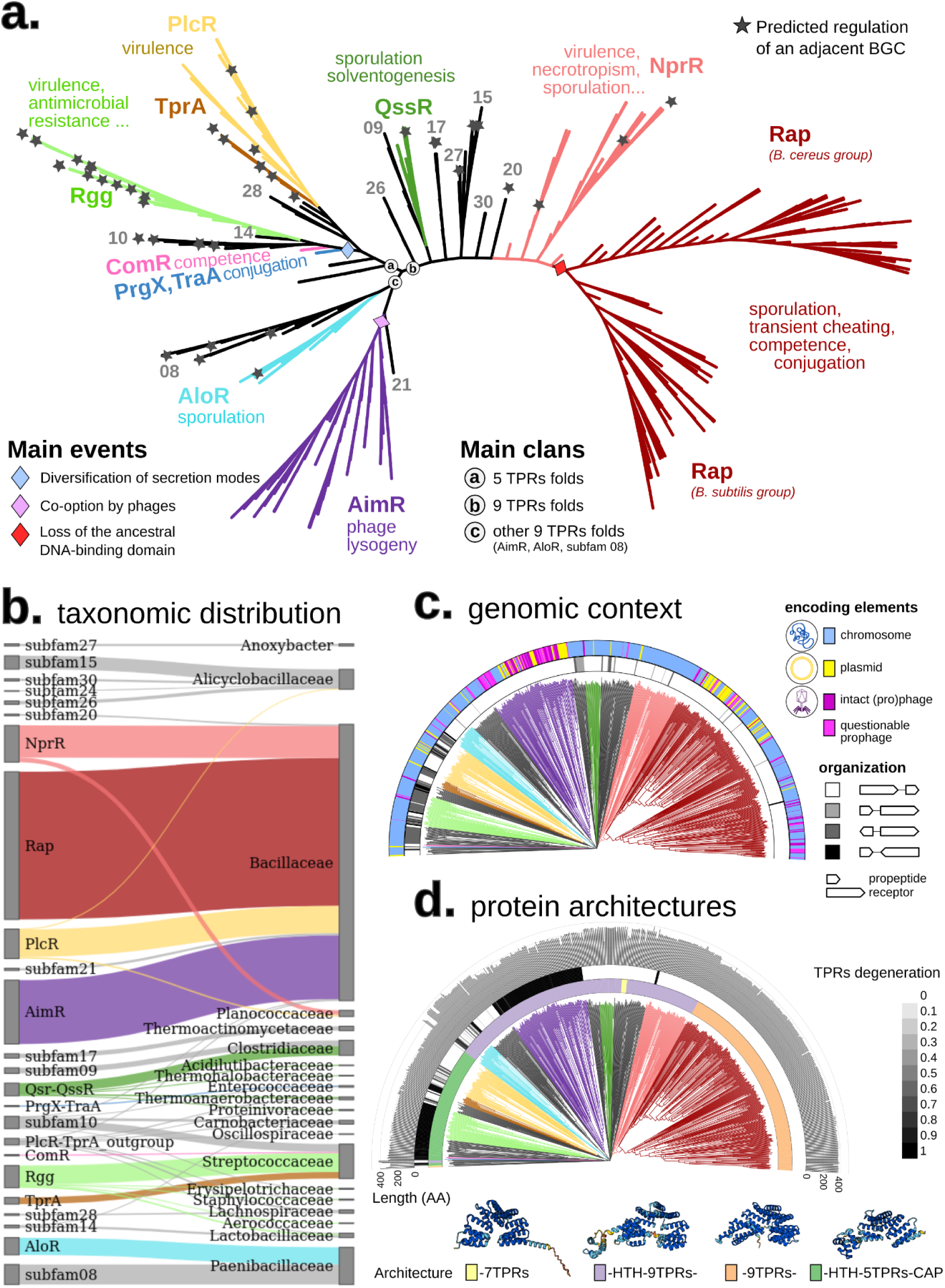
Phylogeny of cytosolic receptors from the RRNPPA family paired with a communication propeptide. **a**) Functional diversity of the RRNPPA family. The tree is unrooted and brench lengths are ignored for visualization purpose (but displayed in panels c and d). Subfamilies with experimental validation of at least one member are highlighted in color. The other subfamilies depicted with a number in grey correspond to high-confidence candidate subfamilies detected with RRNPP_detector in30. Biological processes experimentally shown to be regulated in a density-dependent manner by a QS system are displayed for each validated subfamily. A star mapped to a leaf indicates a predicted regulation of an adjacent biosynthetic gene cluster by the corresponding QS system. **b)** Distribution and prevalence of the different members of each RRNPPA subfamily into the different taxonomic families. **c)** Genomic orientation and encoding element of the receptor - adjacent propeptide pairs. **d)a** The first colorstrip indicates the domain architecture of each receptor. A representative fold for each domain architecture is displayed in the legend (AlphaFold models of subfamily 27, NprR, Rap and PlcR, respectively) with an indication of the implied events from panel a) at the origin of each fold/architecture. The second colorstrip gives the degeneration score of TPR sequences of each receptor (given as 1 - TprPred_likelihood, as in26). The histogram shows the length (in amino-acids) of each receptor

According to Declerck *et al*^26^, receptors with degenerated TPRs correspond to a more diverged state from the common ancestor of RRNPPA receptors than receptors with non-degenerated TPRs. In clan A of 5 TPRs folds, the differential degeneracy levels of TPR sequences suggest that the clade composed of PlcR and TprA receptors have less diverged from the last common ancestor of 5 TPRs folds than the clade composed of PrgX, TraA, ComR and Rgg receptors (**Figure 2d**). While the taxonomic diversity is large in clan A (**Figure 2a-b**), the PrgX-TraA-ComR-Rgg clade is specific to non-spore forming taxa from the *Lactobacillales* order, suggesting indeed a recent innovation. The emergence of the PrgX-TraA-ComR-Rgg clade was accompanied with a diversification in the secretion of the communication peptide. The cognate propeptides of these receptors are either exported through the alternative PptAB translocon (as opposed to the ancestral SEC translocon)^25,28^ or may correspond to leaderless communication peptides involving a yet uncharacterized secretory process^40^.

The degeneracy level of TPRs is lower in receptors from clan B than in receptors from clan C, suggesting that clan B have less diverged than clan C from the last common ancestor of 9 TPRs folds. The clan B encompasses a substantial taxonomic diversity, with taxa distributed in both the *Clostridia* and *Bacilli* classes, (including many extremophiles) (**Table S1**). Its topology implies that the loss of the ancestral HTH DNA-binding domain was the founding event of both the Rap subfamily and the subfamily 17 of *Anoxybacter fermentans*.

The clan C contains both systems from the *Paenibacillaceae* taxonomic family (AloR, subfamily 08) and AimR systems used by temperate phages of *Bacillaceae* to regulate the lysis-lysogeny decision. Consistent with the fact that viruses evolve quickly, the AimR receptors harbor the most degenerated TPR sequences (**Figure 2d**). The phylogeny reconstructed by FoldTree suggests that viral communication systems of the AimR family have been co-opted by bacteriophages from an ancestral receptor that they share with receptors of *Paenibacillaceae*. Consistently, the cognate mature communication peptides of receptors from the AimR subfamily and the subfamily 08 of *Paenibacillaceae* are highly similar, with the widespread presence of the DPG motif across peptides of both subfamilies (**Table S1**).

Whether the last common ancestor of all RRNPPA receptors had 9 TPRs or 5 TPRs still remains an open question. While 9 TPRs folds are mainly associated with dormancy regulation (*e.g.* sporulation in bacteria, lysogeny in phages), 5 TPRs are mostly associated with the density-dependent regulation of biosynthetic gene clusters (**Figure 2a**). Identifying an outgroup with the search of remote structural homologs could be an interesting approach to help deciphering the root and the ancestral state of the family.

## Discussion

Our exploration of the properties of scalable structural phylogenetics based on local alphabets has shown for the first time the viability of these methods on a large scale. The intrinsically slower evolutionary rate of structures allows for the reliable inference of deeper phylogenies where sequence erosion would challenge conventional sequence-based approaches. Viral evolution, quickly evolving extracellular proteins and protein families with histories stretching back to the first self replicating cells are among the many cases that can be revisited with these new techniques. As can be seen from our benchmarking efforts using thousands of families derived using structural or sequence homology, our approach remains robust at varying levels of allowable intrafamily divergence allowing for a broad applicability of structural trees all while remaining capable of resolving deeper relationships. To illustrate the utility of FoldTree, we have chosen to do a deep dive into one such test case, focusing on the fast evolving RRNPPA family of cytosolic communication receptors encoded by Firmicutes bacteria, their conjugative elements and their viruses. The phylogeny reconstructed by FoldTree includes, for the first time, all described RRNPPA subfamilies^30^. Remarkably, despite their significant divergence, the underlying diversifying history of subfamilies is more parsimonious than the sequence phylogeny in terms of taxonomy, functions, and protein architectures (**Supp. Figure 16**).

Declerck *et al.*^26^ also speculated that the level of TPR degeneracy in receptors is a marker of divergence from the last common ancestor of the family. In this respect, root to tips lengths are remarkably uniform throughout the entire RRNPPA structural tree with slight differences being meaningful, as the longest branches correspond to receptors with degenerated TPR sequences (**Figure 2d).** Lastly, the tree inferred by FoldTree systematically is consistent with a scenario of late emergence of clades with degenerated TPRs, as a derived state of an ancestor harboring non-degenerated TPRs (**Figure 2d**). Clades with degenerated TPRs have a narrow taxonomic diversity, which support the hypothesis that they are recent innovations.

Our observations regarding the RRNPPA family only could emerge once their structural homology was used to derive their evolutionary history, painting a coherent picture of their functional diversification. We are far from the first ones to have considered protein structure in an evolutionary light. As early as 1975, Eventoff and Rossmann employed the number of structurally dissimilar residues between pairs of proteins to infer phylogenetic relationships by means of a distance method^41^. This approach has been revisited to infer deep phylogenetic trees and networks using different combinations of dissimilarity measures (*e.g.,* RMSD, Q_score_, Z-score) and inference algorithms^15,42–46^. Conformational sampling has been proposed to assess tree confidence when using this approach^14^. Some models have been developed that mathematically describe the molecular clock in structural evolution^47^ or integrate sequence data with structural information to inform the likelihood of certain substitutions^48^. Other studies have modeled structural evolution as a diffusion process in order to infer evolutionary distances^49^, or incorporated it into a joint sequence-structure model to infer multiple alignments and trees by means of bayesian phylogenetic analysis^50,51^. Despite these efforts, to date the quality of structure-based phylogenetics, especially compared to conventional sequence-based phylogenetics, has remained largely unknown, limiting its use to niche applications.

The extensive empirical assessment on thousands of protein families reported, using three indicators of tree quality, demonstrates the high potential of structure-informed phylogenetics. In addition, our comparison of structurally informed maximum likelihood trees to distance-based trees shows that the simpler model has desirable performance characteristics in empirical benchmarks. The first benchmark, the taxonomic congruence score (TCS), measures agreement of tree topologies with the established taxonomic classification of organisms. Individual gene trees can be expected to deviate substantially from the underlying species tree due to gene duplication, lateral transfer, incomplete lineage sorting, or other phenomena. However, the evolutionary history of the underlying species will still be reflected in many parts of the tree—which is quantified by the TCS. All else being equal, tree inference approaches which tend to result in higher TCS over many protein families can be expected to be more accurate. On this metric, we obtained the best trees using FoldTree, which is based on Foldseek’s structural alphabet, and an alignment procedure combining structural and sequence information. After filtering lower quality structures out of the tree building process, tree quality improved when compared to sequence-only trees (**Supp. Figure 11**), indicating that higher confidence models with accurate structural information provide better phylogenetic signal. The slightly worse tree topologies of maximum likelihood methods using partitioned structure and sequence models on the OMA benchmark compared to normal sequence-based analysis on the TCS benchmark may be due to the model’s presumption of independence of the two partitions when calculating tree likelihood (when, in reality, they represent equivalent positions in the final protein). This may cause discrepancies between the partitions in likelihood calculations. When comparing this method to FoldTree using the second benchmark which also measures topological congruence with the known taxonomy, ASTRAL species tree branch support, (**Supp. Figures 14-1**5) we observe marginally worse results for FoldTree when compared to ML on only the less divergent OMA dataset and marginally better results on the more divergent CATH dataset, indicating that FoldTree may not be the best method to deal with finer grained evolutionary histories and that maximum likelihood may offer more accurate topologies close to the leaves. However, even on the ASTRAL benchmark, 35% of tree topologies produced in the OMA dataset had better quartet support than their ML counterpart, indicating that there may be no ‘one size fits all’ solution even for protein families with homology that is readily identifiable by sequence.

The third benchmarking metric measures adherence to a molecular clock. The idea behind this metric is that the time elapsed between the first instance of a protein at the root of a tree until the sequences observable today at the leaves should be constant for all leaves^52,53^. When considering the root-to-tip variances of the trees, the FoldTree trees adhered more closely to a molecular clock than other structural or sequence trees. We acknowledge that in and of itself, adherence to a molecular clock is a weak indicator of the quality of a tree’s topology. Nevertheless, considering the clear, consistent differences obtained, the slightly higher variance of mean percentage identity values of FoldTree alignments used as inputs for tree building (**Supp. Figures 4-5**) and the superiority of FoldTree when measured with the TCS criterion, the difference in adherence to a clock appears to reflect a meaningful performance increase when comparing FoldTree to the other tree inference methods.

Fold geometry evolves at a slower rate than the underlying sequence mutations^54,55^. Structural distances are therefore less likely to saturate over time, making it possible to align proteins correctly and recover the correct topology deeper in the tree with greater certainty. This could be observed in our results on the distant, structurally defined CATH families. Interestingly, however, FoldTree distinguished itself in TCS comparisons even at divergence times when homology is identifiable using sequence to sequence comparisons and appears only marginally worse when compared with ASTRAL. This may be due to the fact that under conditions where the distance matrix between proteins represents a noisy version of true evolutionary distances between proteins, neighbor joining trees are guaranteed to return the true topology below a certain level of noise^56^. We hypothesize that structural alignment rather than purely sequence based alignment allows for the identification of truly functionally equivalent residues and that the mean rate of replacement across all amino acids of structurally equivalent positions allows for the creation of a distance matrix with lower noise that is a more accurate representation of the evolutionary time elapsed between pairs of proteins. In topologies proposed using maximum likelihood approaches, the likelihood of a given tree is impacted by the substitutions implied by that topology across all sites. These substitutions are not considered in the context of the structure but rather their likelihood is determined by the mean probability of all substitutions across all sites of all proteins considered when constructing the substitution rate matrix being used^57^. At larger evolutionary distances, where cumulated sequence erosion has rendered proteins unrecognizable at the sequence level, these average substitution probabilities coupled to a sequence-based alignment appear to provide a less reliable signal than a simple metric such as the percentage identity after structural alignment.

FoldTree is fine-grained enough to account for small differences between input proteins at shorter divergence times and perform in a manner comparable to state of the art maximum likelihood approaches, overcoming the often mentioned shortcoming of structural phylogenetics. However, its real strength lies in comparison at longer evolutionary distances where its topologies appear more robust than other approaches. The projection of each residue onto a structural character is locally influenced by its neighboring residues rather than global steric changes, Foldseek’s representations of 3D structures are well suited to capture phylogenetic signals when comparing homologous proteins. In contrast, global structural similarity measures, such as the TM score, are confounded by conformational fluctuations which involve steric changes that are much larger in magnitude than the local changes observed between functionally constrained residues during evolution. Moreover, since Foldseek represents 3D structures as strings, the computational speed-ups and techniques associated with string comparisons implemented in MMseqs^58^ are applied to structural comparisons, making the FoldTree pipeline fast and efficient.

Recently the fold universe has been revealed using AlphaFold on the entirety of the sequences in UniProt and the ESM model^11^ on the sequences in MGNIFY^59^ to reach a total of nearly one billion structures. The UniProt structures inferred by AlphaFold have recently been systematically organized into sequence- and structure-based clusters, shedding light on novel fold families and their possible functions^17,60^. In future work it may be desirable to add an evolutionary layer of information to this exploration of the fold space using structural phylogenetics to further refine our understanding of how this extant diversity of folds emerged.

In conclusion, this work shows the potential of structural methods as a powerful tool for inferring evolutionary relationships among proteins. For relatively close proteins, structure-informed tree inference rivals sequence-only inference, and the choice of approach should be tailored to the specific question at hand and the available data. For more distant proteins, structural phylogenetics opens new inroads into studying evolution beyond the “twilight” zone^61^. We believe that there remains much room for improvement in refining phylogenetic methods using the tertiary representation of proteins and hope that this work serves as a starting point for further exploration of deep phylogenies in this new era of AI-generated protein structures.

## Methods

No statistical methods were used to predetermine sample size.

### OMA HOG selection for large scale benchmark

The OMA set of protein families consists of “root hierarchical orthologous groups” (root HOGs) which are derived from all-vs-all sequence comparisons^62^. The quest for orthologs benchmarking dataset^63^ consists of 78 proteomes.The 2020 release of this dataset was used as input into the OMA orthology prediction pipeline^62^ (version 2.4.1). A random selection of at most 500 orthologous groups with at least 10 proteins were compiled for each group of HOGs that were inferred to have emerged in different ancestral taxa (Bacteria, Bilateria, Chordata, Dikarya, Eukaryota, Eumetazoa, Euteleostomi, Fungi, LUCA, Opisthokonta and Tetrapoda). The UniProt identifiers of the proteins within each group were used as input to the FoldTree pipeline.

### CATH family selection for large scale benchmark

CATH structural superfamilies are constructed using structural comparisons and classification^37^. Each level of classification designates a different resolution of structural similarity. These are delineated as Class, Architecture, Topology and Homology. We chose to investigate tree quality using input sets within the same homology classification as well as sets within the same topology. We selected a random subsample of at most 250 proteins (or the number of proteins within the family if there were less) from each family for 635 CATH families. Each CATH family contains the PDB identifiers and chains of the structures they correspond to. We filtered the input lists of proteins in order to only include one representative from each genus as defined by the taxonomic data accompanying each protein in order to avoid redundancy in the dataset.

The PDB files were programmatically obtained from the PDB database. 3D structures of monomers corresponding to the chain identified in the CATH classification for each fold were extracted from PDB crystal structures using Biopython. PDBfixer from the OpenMM^64^ package was used to fix crystal structures with discontinuities, non-standard residues or missing atoms before tree building since these adversely affect structural comparisons.

### Structure tree construction

Sets of homologous structures were downloaded from the AFDB or PDB and prepared according to the OMA and CATH dataset sections above. Foldseek^17^ is then used to perform an all vs all comparison of the structures.

Structural distances between all pairs are compiled into a distance matrix which is used as input to quicktree^65^ to create minimum evolution trees. These trees are then rooted using the MAD method^66^. Foldseek (release: 7-04e0ec8) has two alignment modes where character based structural alignments are performed and are scored using the 3Di substitution matrix or a combination of 3Di and amino-acid substitution matrices. Foldseek was used with the default weighting of amino acids and 3di alignment during alignment scoring. A third mode, using TMalign to perform the initial alignment was not used. It is then possible to output the fraction of identical amino acids from the 3Di and amino acid based alignment (Fident), the LDDT (locally derived using Foldseek’s implementation) score and the TM score (normalized by alignment length). This results in a total of 6 structural comparison methods. We then either directly used the raw score or applied a correction to the scores to transform them to the distance matrices so that pairwise distances would be linearly proportional to time (**Supp. methods**). This resulted in a total of 12 possible Foldseek-based trees for each set of input proteins. To compile these results, Foldseek was used with alignment type 0 and alignment type 2 flags in two separate runs with the ‘--exhaustive-search’ flag. The output was formatted to include lddt and alntmscore columns. The pipeline of comparing structure- and sequence-based trees is outlined in **Supp. Figure 1**.

Before starting the all vs all comparison of the structures we also implemented an optional filtering step to remove poor AlphaFold models with low pLDDT values. If the user activates this option, the pipeline removes structures (and the corresponding sequences) with an average pLDDT score below 40, before establishing the final protein set and running structure and sequence tree building pipelines. We performed similar benchmarking experiments on filtered and unfiltered versions of the OMA dataset to observe the effect of including only high quality models in the analysis.

### Sequence based tree construction

Sets of sequences and their taxonomic lineage information were downloaded using the UniProt API. Clustal Omega (version 1.2.4)^67^ or Muscle5 (version 5.0)^68^ was then used to generate a multiple sequence alignment on default parameters. This alignment was then used with either FastTree(version 2.1)^69^ on default parameters or IQ-TREE (version 1.6.12 using the flags LG+I+G) to generate a phylogenetic tree. Finally, this tree was rooted using the MAD (version 1775932) method on default parameters.

### Taxonomic congruence metric for phylogenetic trees

Taxonomic lineages were retrieved for each sequence and structure of each protein family via the UniProt API. It is assumed that the vast majority of genes will follow an evolutionary trajectory that mirrors the species tree with occasional loss or duplication events. The original development and justification for this score to measure tree quality in an unbiased way can be found in the following work ^35^. In this version of the metric we reward longer lineage sets towards the root by calculating a score for each leaf from the root to the tip. This metric favors correct topology in further back in chronological and evolutionary time, highlighting the utility of incorporating structural information into tree building.

The agreement of the tree with the established taxonomy (from UniProt) can be calculated recursively in a bottom up fashion when traversing the tree using equation 1. Leaves of trees were labeled with sets representing the taxonomic lineages of each sequence before calculating taxonomic congruence. Set elements that shared between all leaves were not included in the calculation of the score since they are uninformative.

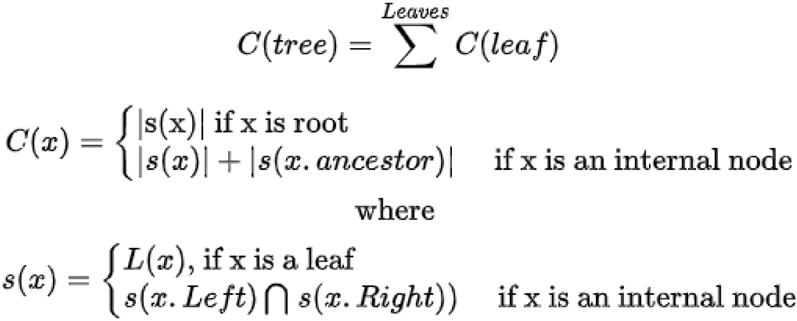

**Equation 1**- taxonomic congruence metric. This score is used to measure the agreement of binary tree topologies with the known species tree. s(x) denotes the set of lineages found in the tree node x. C(x) denotes the congruence score of node x based on its two child nodes. L(x) denotes the labels of leaves. The total score of a tree is defined as the sum of the leaf scores. The code to calculate this metric is available on the git repository. Clades shared between all leaves of a tree are excluded from the calculation as they are uninformative

Both structure and sequence trees were rooted using the MAD method to make TCS comparisons between the methods equivalent. To compare large collections of trees with varying input set sizes, we normalized the congruence scores of trees by the number of the proteins in the tree.

### Molecular clock adherence quantification

Ultrametricity^70^ describes the consistency of tip to root lengths of a given phylogenetic tree. If a tree building approach has an accurate molecular clock on all branches, the amount of inferred evolutionary time elapsed between the root and all of the extant species should be equivalent and proportional to real time. This would imply that the sums of branch length along a lineage from the root to any tip of the tree should be equivalent since the amount of clock time elapsed from the common ancestor until the sequencing of species in the present day is the same.

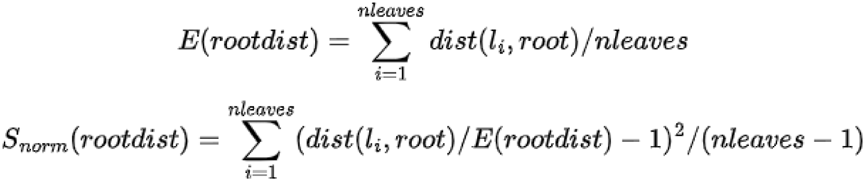

**Equation 2**- To derive a unified metric for adherence to a molecular clock that could easily be applied to the trees generated by different methods, we normalized the branch lengths to center the distribution of root to tip lengths at 1. We then measured the variance of these normalized root to tip lengths. E(.) represents the average root to tip length for a given tree. S_norm_(.) represents the variance of these normalized root to tip distances. dist(l_i_,root) denotes the length of the tip (l_i_) to root.

To describe the degree of adherence to a molecular clock of the different methods of structural tree derivation, we measured the length of root-to-tip distances of a given tree (equation 2). We then normalized this collection of distances by their mean and calculated their variance. We compiled this variance measurement for collections of trees with corresponding input protein sets for all methods used to derive trees and compared their distributions.

### RRNPPA phylogeny

The metadata of “strict” known and candidate RRNPPA QSSs described in the RRNPP_detector paper were fetched from TableS2 in the corresponding Supp. Information^30^. The predicted regulations by QSSs of adjacent BGCs were fetched from TableS5. The propeptide sequences were downloaded from the following Github repository: https://github.com/TeamAIRE/RRNPP_candidate_propeptides_exploration_dataset. The 11,939 receptors listed in TableS2 were downloaded from the NCBI Genbank database, and redundancy was removed by clustering at 95% identity with CD-HIT^71^, yielding 1,418 protein clusters. The Genbank identifiers of the 11,939 receptors were used as queries in the UniProt Retrieve/ID mapping research engine (https://www.uniprot.org/id-mapping) to retrieve corresponding UniProt/AlphaFoldDB identifiers. 768 protein clusters successfully mapped to at least one UniProt/AlphaFoldDB identifier. The 768 predicted protein structures were downloaded and Foldseek was used to perform an all vs all comparison. Based on our benchmarking results we used the Fident scores from a comparison using amino-acid and 3Di alphabet alignment scoring (alignment mode 1 in Foldseek). Since this family had undergone domain architecture modifications, we decided to extract the structural region between the first and last positions of each fold where 80% of all of the other structures in the set mapped. With these core structures we performed a second all vs all comparison (this process is outlined in **Supp. figure 2**). We again used the Fident scores (alignment mode 1) and the statistical correction from **Supp. equation 8** to construct a distance matrix between the core structures. This matrix was then used with FastME^72^ to create a distance based tree. The resulting tree was annotated with ITOL^73^, using the metadata available in **Table S1**. To derive the sequence-based phylogeny, we built a multiple sequence alignment (MSA) of receptors, using mafft^74^ with the parameters –maxiterate 1000 –localpair for high accuracy. The MSA was then trimmed with trimAl^75^ under the -automated 1 mode optimized for maximum likelihood reconstruction. The trimmed alignment of 304 sites was given as input to IQ-TREE^2^ to infer a maximum likelihood phylogenetic under the LG+G model with 1000 ultrafast bootstraps.

## Acknowledgements

We thank the Dessimoz lab members for thoughtful discussions on the topic of structural evolution and their encouragement and input on this work. We also gratefully acknowledge helpful suggestions by Pedro Beltrao.

The work was supported by SNSF grant 216623 to C.D.. M. L. is a recipient of a doctoral scholarship from Agencia Nacional de Investigación e Innovación (ANII), Uruguay.

## Author contributions

David Moi designed and wrote the treebuilding pipeline and analysis pipelines, collected benchmarking data for CATH structural families, carried out large scale analysis for benchmarking, generated trees for protein families, wrote the FoldTree software, and drafted the manuscript. Charles Bernard collected data relevant to the bacterial signaling case study, analyzed and annotated the case study in light of the existing literature and wrote the corresponding sections of the paper. Martin Steinegger contributed advice and feedback on the structural distance measures evaluated in this paper, and contributed to the FoldTree software. Yannis Nevers collected HOG benchmarking data and curated examples of protein families to test the pipeline. Mauricio Langlieb wrote the documentation and collected benchmarking data and curated examples of protein families. Christophe Dessimoz supervised the project and contributed to the conception of the study, the interpretation of results, and the manuscript writing.

## Supplementary Information Guide

1. Supplementary data: The homolog list of RRNPPA sequences and their metadata is available in the RRNPPAlist.xls file. In the text it is referred to as **Table S1**.
2. Supplementary methods, results and discussion are found in the SI section pdf

## Code and Data availability

All UniProt identifiers necessary to replicate the experimental results are available on Zenodo: https://doi.org/10.5281/zenodo.14441021

The FoldTree pipeline is available on GitHub: https://github.com/DessimozLab/fold_tree

The FoldTree Jupyter notebook makes it possible to compute and visualize trees from protein structures in the browser, e.g. using the Google Colab cloud service: https://colab.research.google.com/github/DessimozLab/fold_tree/blob/main/notebooks/FoldTree.ipynb

All metadata used to annotate the RRNPPA phylogeny are available in the supplementary data file or on the Zenodo archive.

## References

1. Kozlov, A. M., Darriba, D., Flouri, T., Morel, B. & Stamatakis, A. RAxML-NG: a fast, scalable and user-friendly tool for maximum likelihood phylogenetic inference. Bioinformatics 35, 4453–4455 (2019).

2. Minh, B. Q., Trifinopoulos, J., Schrempf, D., Schmidt, H. A. & Lanfear, R. IQ-TREE version 2.0: tutorials and Manual Phylogenomic software by maximum likelihood. URL http://www.iqtree.org (2019).

3. Bouckaert, R. et al. BEAST 2.5: An advanced software platform for Bayesian evolutionary analysis. PLoS Comput. Biol. 15, e1006650 (2019).

4. Laumer, C. E. et al. Revisiting metazoan phylogeny with genomic sampling of all phyla. Proc. Biol. Sci. 286, 20190831 (2019).

5. Li, Y., Shen, X.-X., Evans, B., Dunn, C. W. & Rokas, A. Rooting the Animal Tree of Life. Mol. Biol. Evol. 38, 4322–4333 (2021).

6. Schultz, D. T. et al. Ancient gene linkages support ctenophores as sister to other animals. Nature 618, 110–117 (2023).

7. Zea, D. J., Monzon, A. M., Parisi, G. & Marino-Buslje, C. How is structural divergence related to evolutionary information? Mol. Phylogenet. Evol. 127, 859–866 (2018).

8. Lecomte, J. T. J., Vuletich, D. A. & Lesk, A. M. Structural divergence and distant relationships in proteins: evolution of the globins. Curr. Opin. Struct. Biol. 15, 290–301 (2005).

9. Viksna, J. & Gilbert, D. Assessment of the probabilities for evolutionary structural changes in protein folds. Bioinformatics 23, 832–841 (2007).

10. Tunyasuvunakool, K. et al. Highly accurate protein structure prediction for the human proteome. Nature 596, 590–596 (2021).

11. Lin, Z. et al. Evolutionary-scale prediction of atomic-level protein structure with a language model. Science 379, 1123–1130 (2023).

12. Zhang, Y. & Skolnick, J. TM-align: a protein structure alignment algorithm based on the TM-score. Nucleic Acids Res. 33, 2302–2309 (2005).

13. Le, Q., Pollastri, G. & Koehl, P. Structural alphabets for protein structure classification: a comparison study. J. Mol. Biol. 387, 431–450 (2009).

14. Malik, A. J., Poole, A. M. & Allison, J. R. Structural Phylogenetics with Confidence. Mol. Biol. Evol. 37, 2711–2726 (2020).

15. Bujnicki, J. M. Phylogeny of the restriction endonuclease-like superfamily inferred from comparison of protein structures. J. Mol. Evol. 50, 39–44 (2000).

16. Balaji, S. & Srinivasan, N. Use of a database of structural alignments and phylogenetic trees in investigating the relationship between sequence and structural variability among homologous proteins. Protein Eng. 14, 219–226 (2001).

17. van Kempen, M. et al. Fast and accurate protein structure search with Foldseek. Nat. Biotechnol. (2023) doi:10.1038/s41587-023-01773-0.

18. Puente-Lelievre, C. et al. Tertiary-interaction characters enable fast, model-based structural phylogenetics beyond the twilight zone. bioRxiv 2023.12.12.571181 (2023) doi:10.1101/2023.12.12.571181.

19. Xu, J. & Zhang, Y. How significant is a protein structure similarity with TM-score = 0.5? Bioinformatics 26, 889–895 (2010).

20. Mariani, V., Biasini, M., Barbato, A. & Schwede, T. lDDT: a local superposition-free score for comparing protein structures and models using distance difference tests. Bioinformatics 29, 2722–2728 (2013).

21. Clewell, D. B. & Weaver, K. E. Sex pheromones and plasmid transfer in Enterococcus faecalis. Plasmid 21, 175–184 (1989).

22. Rudner, D. Z., LeDeaux, J. R., Ireton, K. & Grossman, A. D. The spo0K locus of Bacillus subtilis is homologous to the oligopeptide permease locus and is required for sporulation and competence. J. Bacteriol. 173, 1388–1398 (1991).

23. Kalamara, M., Spacapan, M., Mandic-Mulec, I. & Stanley-Wall, N. R. Social behaviours by Bacillus subtilis: quorum sensing, kin discrimination and beyond. Mol. Microbiol. 110, 863–878 (2018).

24. Even-Tov, E., Omer Bendori, S., Pollak, S. & Eldar, A. Transient Duplication-Dependent Divergence and Horizontal Transfer Underlie the Evolutionary Dynamics of Bacterial Cell-Cell Signaling. PLoS Biol. 14, e2000330 (2016).

25. Felipe-Ruiz, A., Marina, A. & Rocha, E. P. C. Structural and Genomic Evolution of RRNPPA Systems and Their Pheromone Signaling. MBio 13, e0251422 (2022).

26. Declerck, N. et al. Structure of PlcR: Insights into virulence regulation and evolution of quorum sensing in Gram-positive bacteria. Proc. Natl. Acad. Sci. U. S. A. 104, 18490–18495 (2007).

27. Gallego Del Sol, F., Penadés, J. R. & Marina, A. Deciphering the Molecular Mechanism Underpinning Phage Arbitrium Communication Systems. Mol. Cell 74, 59–72.e3 (2019).

28. Neiditch, M. B., Capodagli, G. C., Prehna, G. & Federle, M. J. Genetic and Structural Analyses of RRNPP Intercellular Peptide Signaling of Gram-Positive Bacteria. Annu. Rev. Genet. 51, 311–333 (2017).

29. Fleuchot, B. et al. Rgg proteins associated with internalized small hydrophobic peptides: a new quorum-sensing mechanism in streptococci. Mol. Microbiol. 80, 1102–1119 (2011).

30. Bernard, C., Li, Y., Lopez, P. & Bapteste, E. Large-Scale Identification of Known and Novel RRNPP Quorum-Sensing Systems by RRNPP_Detector Captures Novel Features of Bacterial, Plasmidic, and Viral Coevolution. Mol. Biol. Evol. 40, (2023).

31. Kotte, A.-K. et al. RRNPP-type quorum sensing affects solvent formation and sporulation in Clostridium acetobutylicum. Microbiology 166, 579–592 (2020).

32. Perez-Pascual, D., Monnet, V. & Gardan, R. Bacterial Cell-Cell Communication in the Host via RRNPP Peptide-Binding Regulators. Front. Microbiol. 7, 706 (2016).

33. Stokar-Avihail, A., Tal, N., Erez, Z., Lopatina, A. & Sorek, R. Widespread Utilization of Peptide Communication in Phages Infecting Soil and Pathogenic Bacteria. Cell Host Microbe 25, 746–755.e5 (2019).

34. Cardoso, P. et al. Rap-Phr Systems from Plasmids pAW63 and pHT8-1 Act Together To Regulate Sporulation in the Bacillus thuringiensis Serovar kurstaki HD73 Strain. Appl. Environ. Microbiol. 86, (2020).

35. Tan, G., Gil, M., Löytynoja, A. P., Goldman, N. & Dessimoz, C. Simple chained guide trees give poorer multiple sequence alignments than inferred trees in simulation and phylogenetic benchmarks. Proc. Natl. Acad. Sci. U. S. A. 112, E99–100 (2015).

36. Van Etten, J. & Bhattacharya, D. Horizontal Gene Transfer in Eukaryotes: Not if, but How Much? Trends Genet. 36, 915–925 (2020).

37. Knudsen, M. & Wiuf, C. The CATH database. Hum. Genomics 4, 207–212 (2010).

38. Mutti, G., Ocaña-Pallarés, E. & Gabaldón, T. Newly developed structure-based methods do not outperform standard sequence-based methods for large-scale phylogenomics. bioRxiv (2024) doi:10.1101/2024.08.02.606352.

39. Bereg, S. & Zhang, Y. Phylogenetic networks based on the molecular clock hypothesis. in Fifth IEEE Symposium on Bioinformatics and Bioengineering (BIBE’05) 320–323 (2005). doi:10.1109/BIBE.2005.46.

40. Aggarwal, S. et al. The leaderless communication peptide (LCP) class of quorum-sensing peptides is broadly distributed among Firmicutes. Nat. Commun. 14, 5947 (2023).

41. Eventoff, W. & Rossmann, M. G. The evolution of dehydrogenases and kinases. CRC Crit. Rev. Biochem. 3, 111–140 (1975).

42. Johnson, M. S., Sali, A. & Blundell, T. L. Phylogenetic relationships from three-dimensional protein structures. Methods Enzymol. 183, 670–690 (1990).

43. Garau, G., Di Guilmi, A. M. & Hall, B. G. Structure-based phylogeny of the metallo-beta-lactamases. Antimicrob. Agents Chemother. 49, 2778–2784 (2005).

44. Lundin, D., Berggren, G., Logan, D. T. & Sjöberg, B.-M. The origin and evolution of ribonucleotide reduction. Life 5, 604–636 (2015).

45. Moi, D. et al. Discovery of archaeal fusexins homologous to eukaryotic HAP2/GCS1 gamete fusion proteins. Nat. Commun. 13, 3880 (2022).

46. Lakshmi, B., Mishra, M., Srinivasan, N. & Archunan, G. Structure-Based Phylogenetic Analysis of the Lipocalin Superfamily. PLoS One 10, e0135507 (2015).

47. Pascual-García, A., Arenas, M. & Bastolla, U. The Molecular Clock in the Evolution of Protein Structures. Syst. Biol. 68, 987–1002 (2019).

48. Arenas, M., Sánchez-Cobos, A. & Bastolla, U. Maximum-Likelihood Phylogenetic Inference with Selection on Protein Folding Stability. Mol. Biol. Evol. 32, 2195–2207 (2015).

49. Grishin, N. V. Estimation of evolutionary distances from protein spatial structures. J. Mol. Evol. 45, 359–369 (1997).

50. Challis, C. J. & Schmidler, S. C. A stochastic evolutionary model for protein structure alignment and phylogeny. Mol. Biol. Evol. 29, 3575–3587 (2012).

51. Herman, J. L., Challis, C. J., Novák, Á., Hein, J. & Schmidler, S. C. Simultaneous Bayesian estimation of alignment and phylogeny under a joint model of protein sequence and structure. Mol. Biol. Evol. 31, 2251–2266 (2014).

52. Smith, S. A., Brown, J. W. & Walker, J. F. So many genes, so little time: A practical approach to divergence-time estimation in the genomic era. PLoS One 13, e0197433 (2018).

53. Wertheim, J. O., Sanderson, M. J., Worobey, M. & Bjork, A. Relaxed molecular clocks, the bias-variance trade-off, and the quality of phylogenetic inference. Syst. Biol. 59, 1–8 (2010).

54. Illergård, K., Ardell, D. H. & Elofsson, A. Structure is three to ten times more conserved than sequence--a study of structural response in protein cores. Proteins 77, 499–508 (2009).

55. Chothia, C. & Lesk, A. M. The relation between the divergence of sequence and structure in proteins. EMBO J. 5, 823–826 (1986).

56. Atteson, K. The performance of neighbor-joining methods of phylogenetic reconstruction. Algorithmica 25, 251–278 (1999).

57. Le, S. Q. & Gascuel, O. An improved general amino acid replacement matrix. Mol. Biol. Evol. 25, 1307–1320 (2008).

58. Steinegger, M. & Söding, J. MMseqs2 enables sensitive protein sequence searching for the analysis of massive data sets. Nat. Biotechnol. 35, 1026–1028 (2017).

59. Richardson, L. et al. MGnify: the microbiome sequence data analysis resource in 2023. Nucleic Acids Res. 51, D753–D759 (2023).

60. Durairaj, J. et al. Uncovering new families and folds in the natural protein universe. Nature (2023) doi:10.1038/s41586-023-06622-3.

61. Rost, B. Twilight zone of protein sequence alignments. Protein Eng. 12, 85–94 (1999).

62. Altenhoff, A. M., et al. OMA standalone: orthology inference among public and custom genomes and transcriptomes. Genome Res. 29, 1152–1163 (2019).

63. Altenhoff, A. M., et al. The Quest for Orthologs benchmark service and consensus calls in 2020. Nucleic Acids Res. 48, W538–W545 (2020).

64. Eastman, P. et al. OpenMM 7: Rapid development of high performance algorithms for molecular dynamics. PLoS Comput. Biol. 13, e1005659 (2017).

65. Howe, K., Bateman, A. & Durbin, R. QuickTree: building huge Neighbour-Joining trees of protein sequences. Bioinformatics 18, 1546–1547 (2002).

66. Tria, F. D. K., Landan, G. & Dagan, T. Phylogenetic rooting using minimal ancestor deviation. Nat Ecol Evol 1, 193 (2017).

67. Sievers, F. & Higgins, D. G. Clustal Omega, accurate alignment of very large numbers of sequences. Methods Mol. Biol. 1079, 105–116 (2014).

68. Edgar, R. C. MUSCLE: multiple sequence alignment with high accuracy and high throughput. Nucleic Acids Res. 32, 1792–1797 (2004).

69. Price, M. N., Dehal, P. S. & Arkin, A. P. FastTree: Computing Large Minimum Evolution Trees with Profiles instead of a Distance Matrix. Mol. Biol. Evol. 26, 1641–1650 (2009).

70. Moore, N. C. A. & Prosser, P. The Ultrametric Constraint and its Application to Phylogenetics. arXiv [cs.AI] (2014).

71. Li, W. & Godzik, A. Cd-hit: a fast program for clustering and comparing large sets of protein or nucleotide sequences. Bioinformatics 22, 1658–1659 (2006).

72. Lefort, V., Desper, R. & Gascuel, O. FastME 2.0: A Comprehensive, Accurate, and Fast Distance-Based Phylogeny Inference Program. Mol. Biol. Evol. 32, 2798–2800 (2015).

73. Letunic, I. & Bork, P. Interactive Tree Of Life (iTOL) v5: an online tool for phylogenetic tree display and annotation. Nucleic Acids Res. 49, W293–W296 (2021).

74. Katoh, K. & Standley, D. M. MAFFT multiple sequence alignment software version 7: improvements in performance and usability. Mol. Biol. Evol. 30, 772–780 (2013).

75. Capella-Gutierrez, S., Silla-Martinez, J. M. & Gabaldon, T. trimAl: a tool for automated alignment trimming in large-scale phylogenetic analyses. Bioinformatics vol. 25 1972–1973 Preprint at 10.1093/bioinformatics/btp348 (2009).

