## supplementary_information for "Structural phylogenetics unravels the evolutionary diversification of communication systems in gram-positive bacteria and their viruses"

David Moi et al.

### Pipeline description

In order to efficiently generate large numbers of phylogenetic trees using structure and sequence based methods for comparative analysis, we built a Snakemake pipeline which is available on the github associated with this project. The pipeline steps are outlined below in Supplementary figure 1.

The different steps of the pipeline are also usable through the python library associated with the project, either through scripting or through the command line interface. Instructions on using the pipeline are available on the project github (https://github.com/DessimozLab/fold_tree). We also show the corecut pipeline used in the construction of the RRNPPA tree. This process ensures that only the domains shared between a large enough proportion of the input set of proteins is use to construct the tree. C and N terminal domains that are outside of this consensus region are then clustered and cluster labels are used to annotate the phylogeny, showing where architecture changes may have happened. The process is schematized below in **Supplementary Figure 3**.

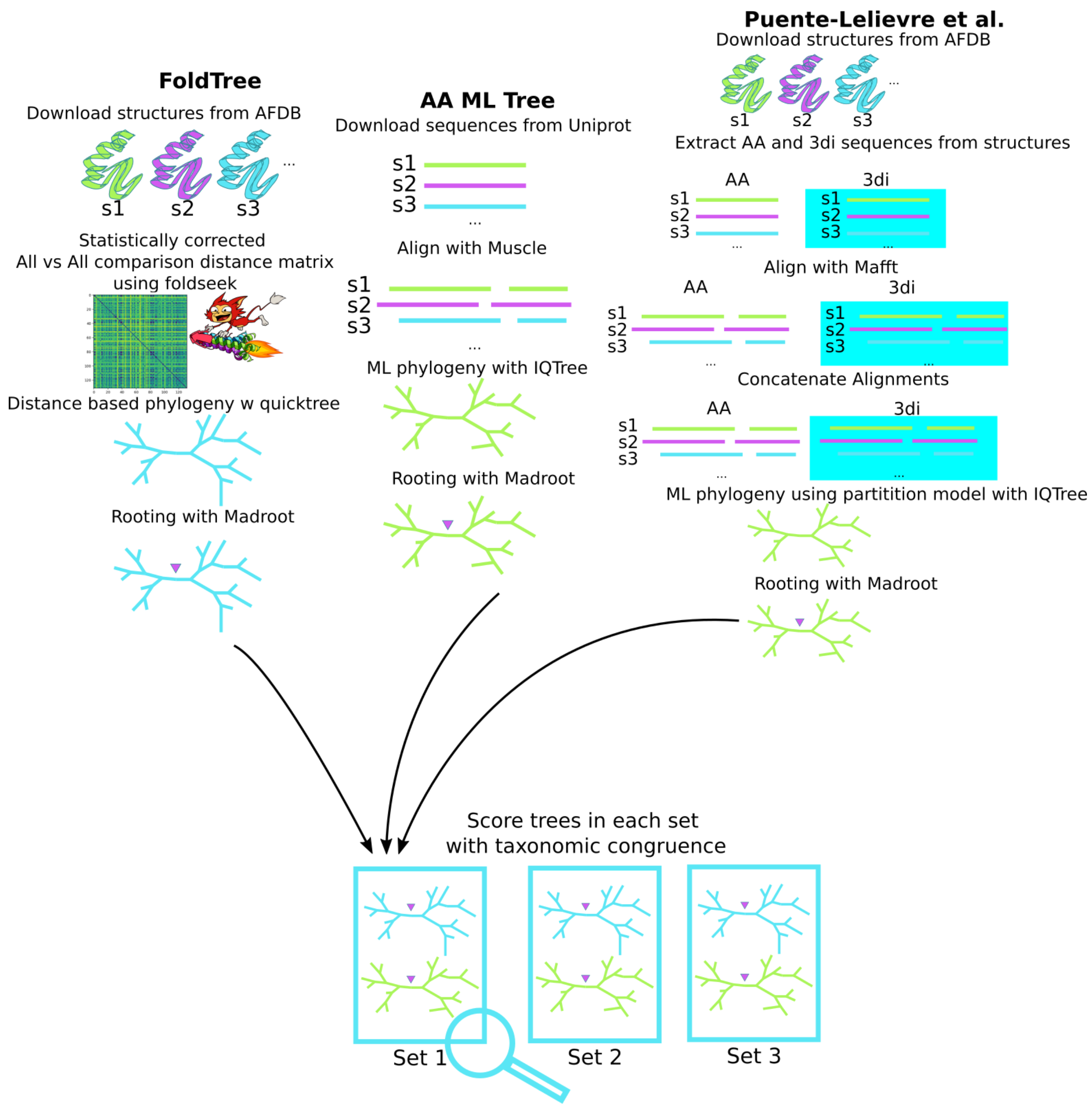


**Supplementary Figure 1** Benchmarking pipeline schematic - Trees are created from equivalent protein sets for structure and sequence trees. On the structural side of the pipeline, all vs all comparisons using foldseek are compiled into a statistically corrected distance matrix. These are used as input for quicktree and rooted with MAD. On the sequence side of the pipeline, the sequences are aligned with Muscle. A maximum likelihood tree is derived using IQTree and then rooted with MAD. The method described by Puente-Lelievre et al. is implemented as reported in their manuscript (Puente-Lelievre et al.). 3di and amino acid characters are aligned using Mafft and then concatenated into an alignment containing both with the same pattern of gaps in both partitions. IQTree is then used with a partition model using the LG and 3di substitution matrices on their respective partitions. Both structural and sequence-based trees are scored using the TCS metric described in *Methods* and outlined below in Supplementary Figure 2.

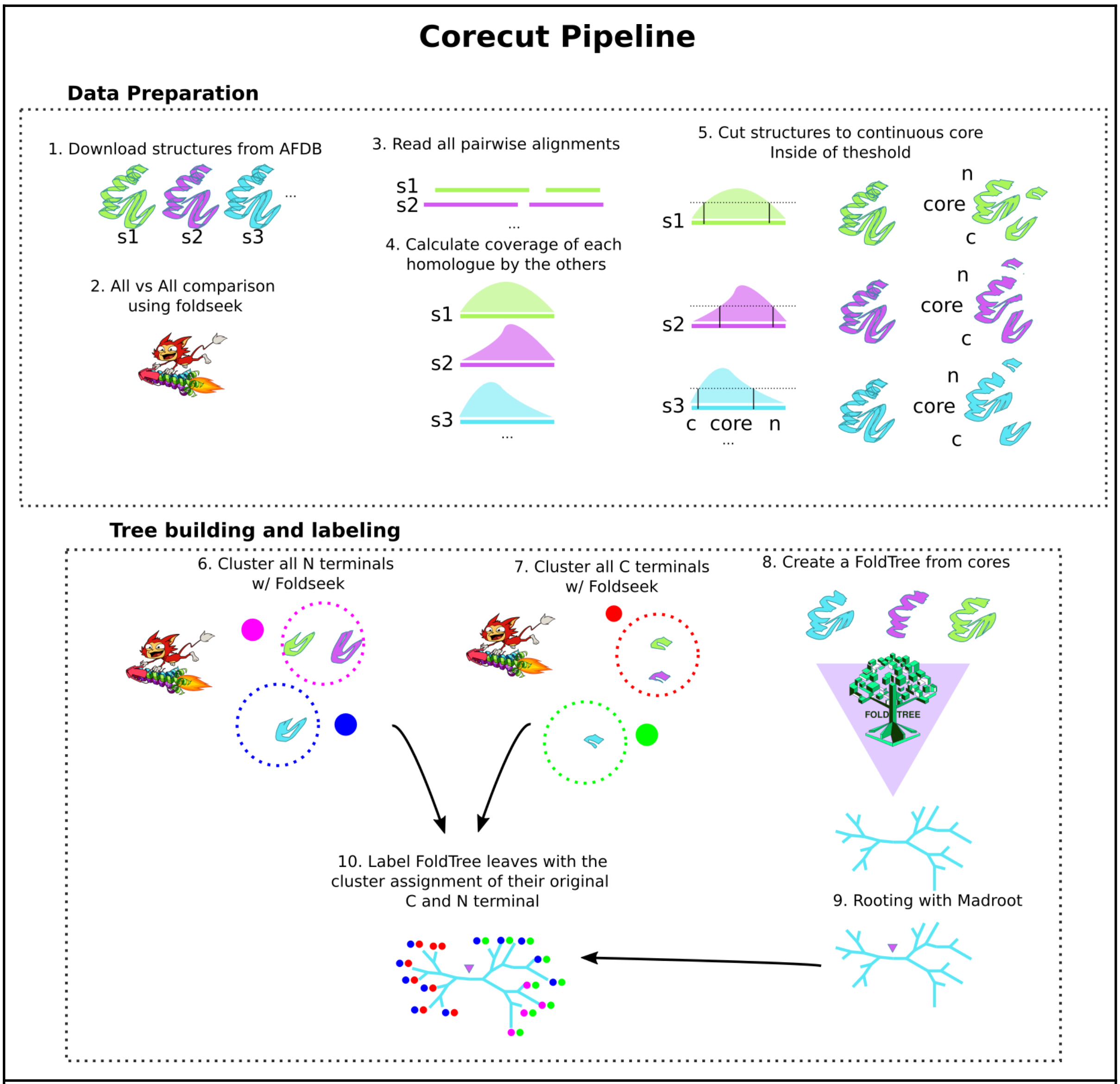


**Supplementary Figure 2** Core Cut pipeline schematic. In steps 1-5 we outline how we used Foldseek to derive a consensus core structure between a group of homologs. This process divides the structure into 3 regions. In the second part of the pipeline ( steps 6-10 ) we show how we constructed clusters for the N and C terminals as well as a FoldTree from the consensus core of the set of homologs. The final output of this pipeline is a FoldTree of the core labeled with the assigned clusters of the C and N terminal of the input structures.

In **Eq. 1** of the main text, we show how we calculate the taxonomic congruence score as a stand-in for tree quality. In order to provide a more intuitive way of representing how this score is derived, we have also included **Supplementary Figure 3** showing a diagram of how the trees are scored recursively using sets to represent lineages.

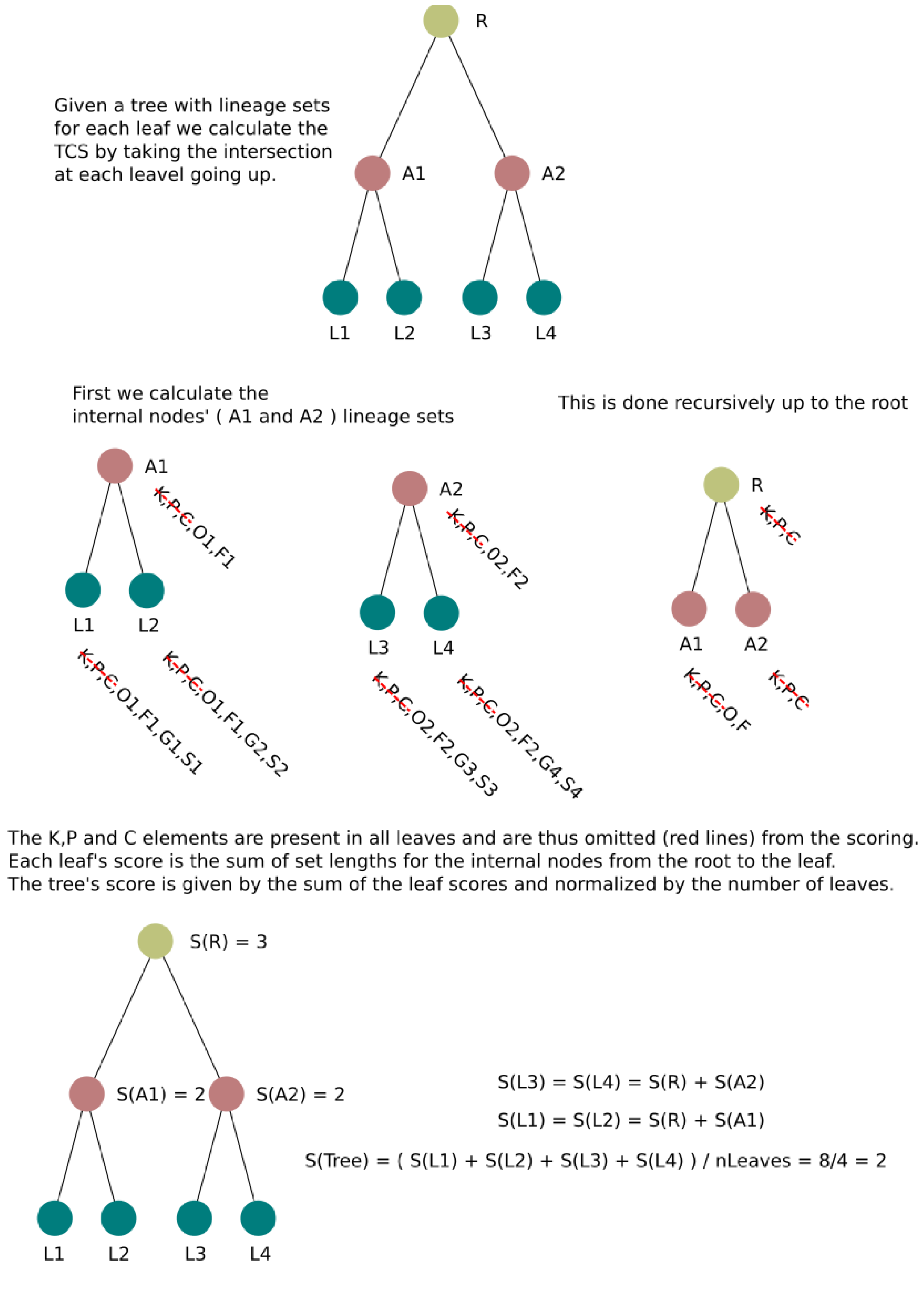


**Supplementary Figure 3 schema of TCS score calculation**
In the example tree shown each leaf is labeled with simple lineage information containing its Kingdom, Phylum, Class, Order, Family, Genus and Species. At each level of the tree,

the sets of internal nodes and the root are determined by the intersection of the two nodes found below. The leaf scores are then calculated by taking the sum of scores of internal nodes from the root to a leaf. The tree score is the sum of these scores normalized by the number of leaves.

The scores presented in the figures comparing methods are normalized by the number of leaves. The scores are impacted by the 'density' of taxonomic information in each clade rather than the size of the trees being considered. It is therefore only correct to consider the comparison of TCS scores within a given family with the same input set of proteins and not between families. Even if two families were to have the same number of proteins from the same species, the true topology of the tree explaining their evolution may have an intrinsically lower TCS score than another family due to the myriad of evolutionary phenomena that do not follow the typical vertical mode of inheritance.

#### Pipeline description

#### Derving MSAs for use with partition model

Three versions of maximum likelihood approaches were tried using the amino acid and 3di representations of the proteins in each of the CATH and OMA families used for benchmarking. The first and second versions are based on using the alignment of either the amino acid or 3di sequences to align the other character set. Thus, the alignment logic used is based on a single alphabet and is transferred over to the other. For benchmarking purposes, we used Mafft on default parameters to align the amino acid sequences and the Mafft textalign option with the 3di substitution matrix used in foldseek to align 3di strings. The gap pattern was then transferred from the amino acid MSA to the 3di strings or from the 3di MSA to the amino acid sequences. This method and its performance on one case study protein family were first described in the following manuscript (Puente-Lelievre et al.).

#### Studying induced pairs from MSAs

Using the MSAs derived from structural and sequence alignments described above, we calculated the pairwise percent identity of the amino acid sequences. These were then used to create distance matrices and calculate distance-based trees in order to investigate the effect of tree building methods on TCS and molecular clock adherence. A statistical correction was applied to the distances derived identically to the correction used in the FoldTree method. This correction is detailed below in **equation 8**. The uncorrected distance matrices were used to calculate the distributions of pairwise identities for each family using different alignment approaches. The summary statistics of these distributions were collected and we then derived the distributions of the mean, variance and skewness for all alignment methods across all families. The results are shown below in **Supp Figure 3-4**.

#### Structure trimming to find the 'core' region

Structural families will often contain representatives that have extra domains not found in the majority of the members of the family. This phenomenon is especially prevalent in prokaryotes where domain shuffling events are common. To find the conserved core of the structure set being compared and only use the phylogenetic signal from this region, we implemented a post processing step after the first all vs. all comparison using Foldseek. The results of the comparisons within a set are mapped to the corresponding amino acid positions of a protein. A continuous region between the first and the last position with over 80% of the dataset mapping to them is designated as the core. The C and N terminals of the protein that are not within this core are removed and a new pdb file is generated with only the core using Biopython's structure module. The set of 'core' proteins can then be used as input to FoldTree. The RRNPPA phylogeny was processed using this technique in order to reduce the input set of proteins to their 'core' before using them as input for tree building. The trimmed regions were clustered and used to label the tree to verify that architecture changes were distributed on the tree parsimoniously.

#### Evolutionary distance correction of structurally derived percent identity or distance based scores

To transform the Fident, LDDT or TM scores into unbiased estimators of evolutionary distances (Rzhetsky and Nei) that are proportional to time, we transformed the raw scores using a model based on a process of random replacement.

In the case where a discrete process changes the characters in the sequence of a protein represented by the structural alphabet over time, we can either estimate the divergence time using the following formula which uses a taylor expansion of the logarithm for numerical stability (Tajima, “Unbiased Estimation of Evolutionary Distance between Nucleotide Sequences”). This formula is applicable for the Fident metric which represents the amount of identical residues aligned with the 3Di alphabet representation of a structure (van Kempen et al.).

**Equ. 1.**

$$\hat{d} = \sum_{i=1}^{k} k^i / in^i$$

where $\hat{d}$ is the divergence time. The parameters k and n denote the number of non-identical residues and length of the sequence, respectively.

In the continuous case where we propose to model the process of two ancestral structures diverging along a bifurcation in the tree. The ancestral structure is present at time 0 and we consider the structural distance metric between its two descents at time t. We model this process of mutation accumulation as a continuous time stochastic process where the logarithm of the randomly varying quantity (the structural distance) follows a pattern composed of brownian motion (noise) and drift (selection) (Øksendal). With first and second terms representing the drift and brownian motion respectively.

$$dS_t = \mu dS_t dt + \sigma S_t W_t$$

**Equ. 2.**

In this case we can write the expected distance between a pair of structures after a time t (in our case the TM or LDDT distances) after diverging from a single ancestral structure. The left hand side of equation 2 represents a small delta in this quantity. Since the expected value of the Wiener process in equation 2 is 0, the expected value after a time t is shown below in Equation 3.

$$E(S_t) = S_0 e^{\mu t}$$

**Equ. 3.**

By admitting that at state $S_0$, after no divergence time, represents a pair of structures that would be identical, then the LDDT and TM scores between a structure and itself will be equal to 1 (and thus $\ln(S_0)=0$). We can take the logarithm of both sides of the expression to give us:

$$ln(E(S_t)) = \mu t$$

**Equ. 4.**

Since $S_t$ must be smaller than $S_0$ for the TM score and LDDT, μ should be negative. Therefore we take the value

$$-ln(E(S_t)) = ct$$

**Equ. 5.**

where c is a positive constant, making the left hand quantity linear in time. In order to calculate this quantity, we use the Taylor expansion of the natural logarithm of this formula as well as in Equation 1. With the following Taylor expansion that converges for all x with an absolute value less than one (Tajima, "Unbiased Estimation of Evolutionary Distance between Nucleotide Sequences").

**Equ. 6.**

$$ln(1 - x) = -\sum_{n=1}^{\infty}(x)^n/n$$

by substituting *1-x = score* we have:

**Equ. 7.**

$$-ln(score) = \sum_{n=1}^{\infty}(1 - score)^n/n$$

In the implementation of this score we only use the first 100 terms of the expansion. Similar to sequence identity, the logarithm of structural similarity scores should be linearly dependent on the drift rate and time. Thus, our proposed method to correct the evolutionary distance is the right-hand side of Equation 7.

However, this does not presuppose that all mutation or drift rates are the same across all evolutionary trajectories (i.e. that the molecular clock is identical on all branches) which may impact minimum evolution tree building approaches (Tajima, "Simple Methods for Testing the Molecular Evolutionary Clock Hypothesis").

In the case of sequence identity using the amino acid alphabet, we use Tajima's correction including the term accounting for random back mutations (Tajima, "Unbiased

Estimation of Evolutionary Distance between Nucleotide Sequences”). We have set this term to b=.93 which would correspond to a saturation of mutations over signal at 7% sequence identity. We also use 100 iterations of the expansion for approximating this score. This score was used to derive an evolutionary distance from the Fident metric reported by Foldseek to create the FoldTree metric as well as correcting the induced pairs of MSAs when exploring the properties of distance based trees without structural alignment.

**Equ. 8.**

$$-bln(1-(p/b)) = \sum_{i}^{k} \frac{k^i}{ib^{i-1}n^i}$$

#### ASTRAL benchmark of quartet support.

The ASTRAL-PRO tool is designed to study the support that an inferred phylogeny gives the leaf quartets that are present in a species tree topology or vice versa ( e.g. deriving a species tree given the quartets present in a collection of inferred phylogenies ). This benchmark was proposed in (Mutti et al.) in order to examine structural vs sequence tree quality compared to the species tree. We ran this benchmark as a complement to our TCS-based topology scoring in order to provide another measure of tree quality that is more sensitive to topological errors closer to the leaves. A complete description of the ASTRAL-PRO algorithm is available in the original manuscript (Zhang and Mirarab).

For each family we used the NCBI taxonomy topology spanning all species included in the input set of proteins using the ete3 ncbi taxonomy package as the species tree to compare to. We then ran ASTRAL-PRO3 from the ASTER package using the -C -T -E flags using the species tree as a reference tree and the phylogeny being scored as the input. In addition to providing support values for species tree branches, also reports implied loss and duplication events when species contain multiple genes within a family. The number of implied loss and duplications are reported for the OMA families used in our benchmarking. Since the CATH dataset’s families were filtered to only include one member of each genus in order to avoid redundancy, only support values for the species tree are compared.

In order to resolve the multi furcations in the NCBI species tree topologies we used ETE3 to resolve them into a binary tree randomly. The resolved input species trees were kept constant for all input gene trees.

#### Alignment properties for families aligned with 3di, Amino Acids or a combination of both.

After deriving MSAs of proteins in each family to construct maximum likelihood trees as well as pairwise comparisons of all families with Foldseek to construct distance trees, we observed the qualities of the distributions of pairwise identities produced by each approach. For each protein family we compiled the mean, variance and skewness of pairwise identities of all pairs in the family. The distributions for each of these values is shown below for the OMA and CATH datasets. By using both 3di and amino acid characters, the mean pairwise identity is increased.

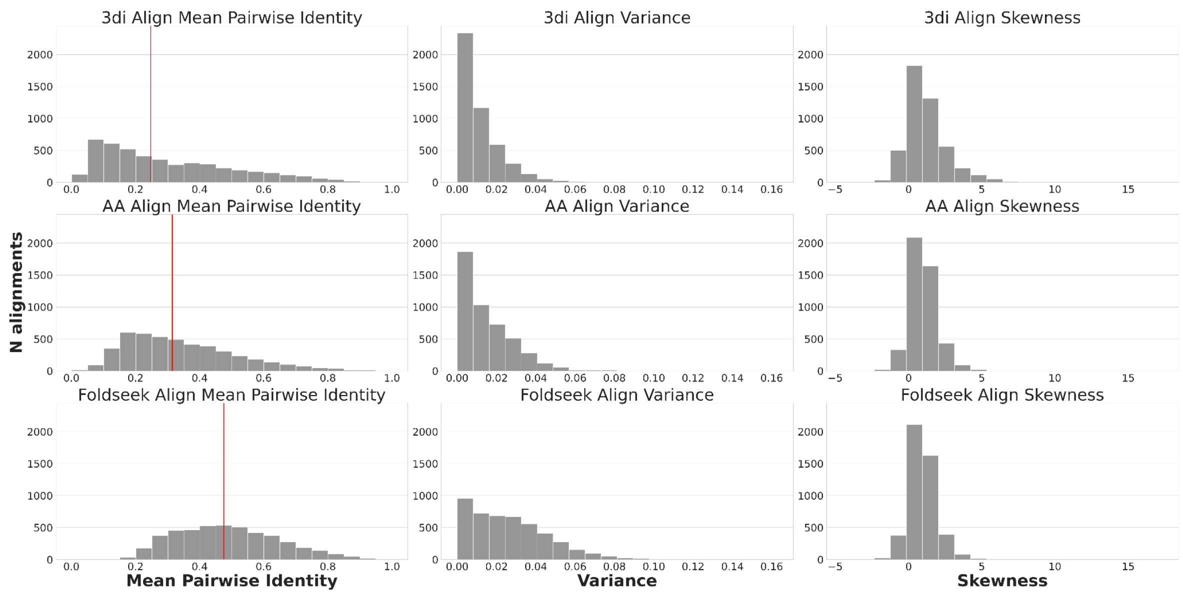


**Supplementary Figure 4 Alignment pairwise identity distributions aligning with amino acids, 3di or both for the OMA dataset** Distributions of mean pairwise identities, variances and skewnesses are shown for all families in the OMA dataset after alignment with either 3di and amino acids ( Foldseek ) or with just one of the two. We observe an increase of mean pairwise identity after aligning with Foldseek accompanied by a higher variance in pairwise identities.

If this increase in pairwise identities reflects the increased detection of equivalent residues across different extant homologs of a protein family, this may reflect a higher quality of input data into tree building. This in turn would positively impact the results observed in the TCS and molecular clock adherence benchmarks. As shown below in **Supp Figure 4**, when the maximum divergence allowed within a protein family is increased, the quality of alignments based exclusively off of amino acids or 3di suffer while using a combination of both again drives up the mean pairwise identity of alignments ( although their percentage identity is still lower than the alignments observed in the OMA dataset ). At lower evolutionary distances, the small differences in alignment may be negligible in their effect on tree topology and molecular clock adherence but as more divergent families enter the set of families analyzed, using structural information brings a significant advantage as seen in **Figure 1** and **Supp Figure 12**.

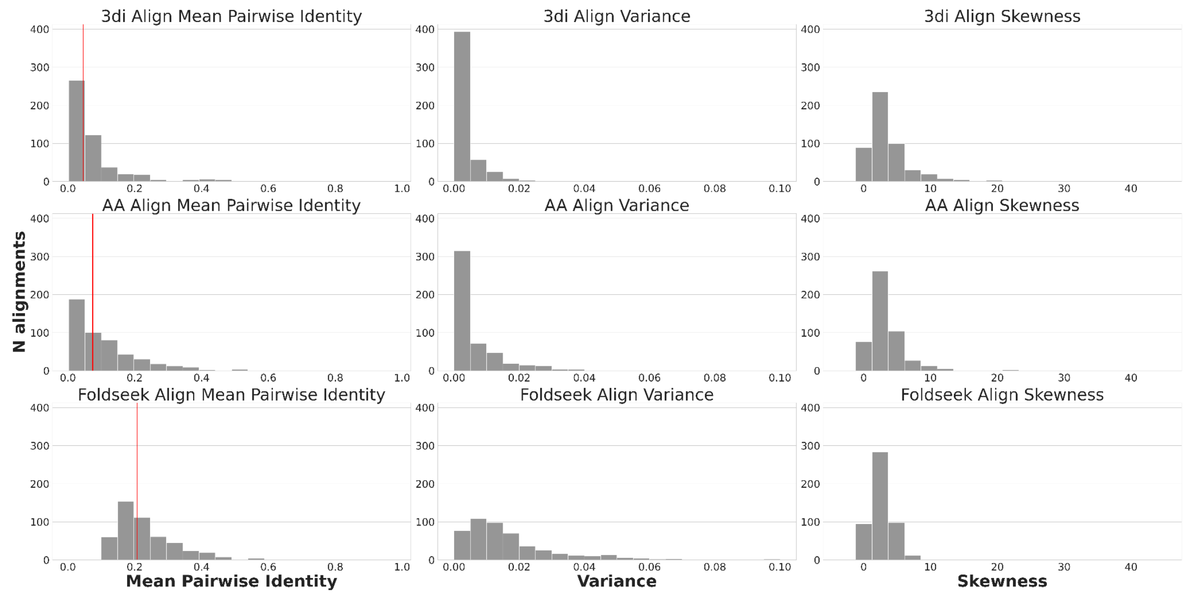


**Supplementary Figure 5 Alignment pairwise identity distributions aligning with amino acids, 3di or both for the CATH dataset** Aggregate distributions of the mean, variance and skewness distributions are shown for percentage identity distributions of all alignments in the CATH dataset. We observe an even larger discrepancy between the mean of pairwise identities than in **supp figure 3** when aligning with just one character set as opposed to both 3di and amino acid in tandem.

In light of this, we split the OMA and CATH datasets into quartiles by alignment pairwise identities. We observe an increase in the difference measured between tree topologies alongside and increase in alignment divergence. These results are shown below in **Supp Figure 12**.

In addition to alignment quality, structural features such as the predicted model quality or pLDDT are important for inferring the relationships between residues and assigning the correct 3di character to each position of the protein. The availability of good models for a protein family of interest should be considered before using a structural phylogenetics approach. To verify this we filtered the OMA dataset to input protein sets whose average AlphaFold model pLDDT greater than 40 and with an overall minimum greater than 30. Families not meeting this criteria were excluded from the analysis. The difference in topology quality over sequence-based trees for the FoldTree metric is shown below in **Supp Figure 10**.

#### Supplementary TCS comparisons

When investigating which type of structural distance metric would be the most appropriate for creating structural trees we considered all of the alignment modes and outputs provided by foldseek. As we observed, not all of them provide an informative signal for the task at hand. Here we present TCS comparisons of trees derived using alignment mode 0 in (only using the structural alphabet) in Foldseek against sequence-based phylogenetics below in **Supp Figure 6**.

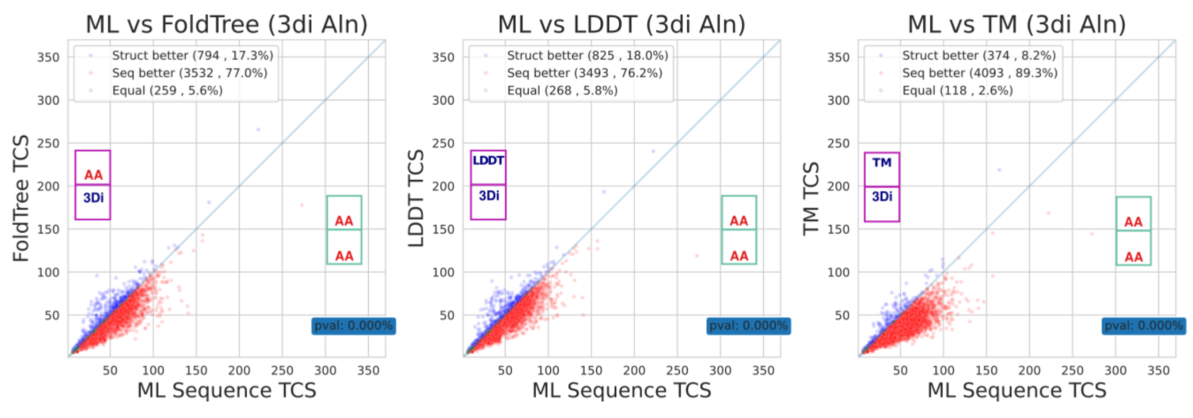


**Supplementary Figure 6.** Structural Distance metrics using only structural character alignment score poorly in terms of TCS on the unfiltered OMA dataset. The P-value for the Wilcoxon ranked test between the TCS scores of structure trees and their corresponding maximum likelihood trees is shown in the blue rectangle in the bottom right. With all structural distance approaches, aligning with only 3di characters produces inferior trees.

We also studied the effect of using statistical corrections on the input distances. In **equations 1-8** we propose models to compensate for back mutations or, in the case of geometric distances, fluctuations in protein geometry which may bring proteins back towards an initial state. After building trees with both corrected and uncorrected measures, we applied both TCS and root to tip variance benchmarks in order to determine if the corrections had impacted either measure of tree quality. In **Supp Figure 7** below, the results of comparing root to tip variance of the corrected and uncorrected distance trees is shown. All results in this manuscript pertaining to tree quality or the presented RRNPPA family are shown for corrected trees, apart from **Supp Figures 7-8**, showing the effects of this correction on molecular clock adherence and TCS respectively.

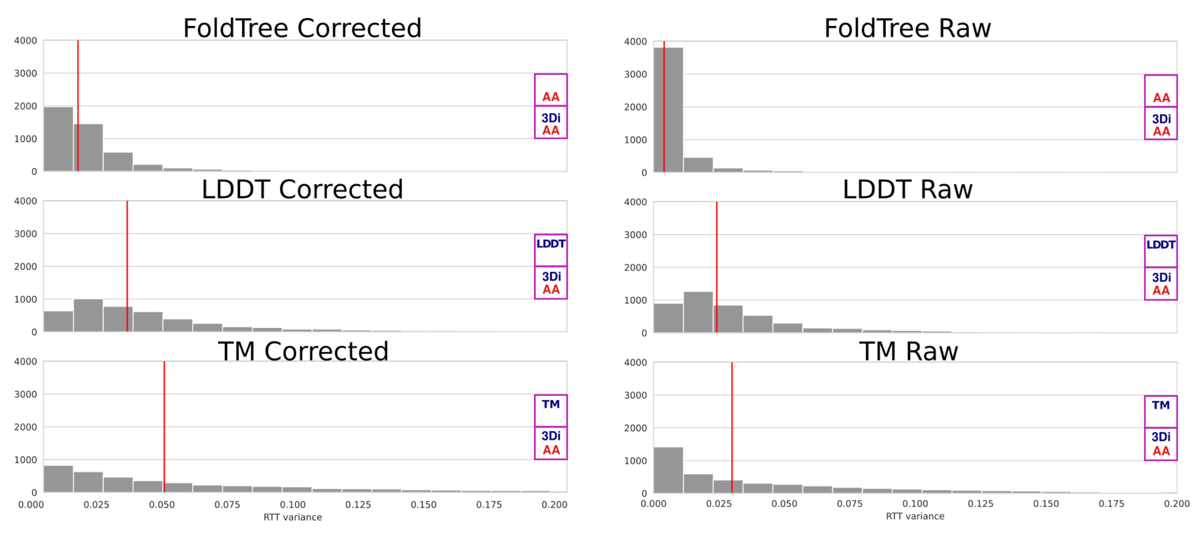


**Supplementary Figure 7. Effect of statistical corrections on molecular clock adherence.** Structural distances or the fraction of identical residues in the case of FoldTree can be used to build trees without modification or statistically corrected using **supp. equations 1-9** before treebuilding. Root to tip variance distributions presented in **Figure 1** are the statistically corrected versions of all distance trees. The uncorrected distances may saturate at a maximum finite distance. After statistical correction shown in **equation 9** the evolutionary distance reported by FoldTree will go towards an infinite value when sequence identity approaches 7% and the statistically corrected LDDT and TM distances will approach an infinite value when measured structural similarity approaches 0. The corrected score used by FoldTree retains a lower root to tip variance than maximum likelihood approaches and the other structural distance metrics before and after correction as seen in **Figure 1**. The median value of each distribution is shown with a red vertical line.

While the trees using corrected distance metrics appear less clock-like, they still retain a lower root to tip variance than typical sequence-based approaches as shown in **Figure 1** and below in **Supp Figure 12**. We consider the corrected distances to be more realistic than the raw distances since they have the property of tending towards infinity when the comparison of two proteins shows a signal that is below the expected value of the comparison of two random proteins.

When considering differences in TCS across the corrected and uncorrected structural distances we experimented with, it appears that the effect of statistical correction is negligible as shown below in **Supp Figure 8**. We believe taking into account mutations back to an original state ( or geometric fluctuations back to an original state in the case of geometric distances ) they represent a more realistic tree in terms of topology and branch length.

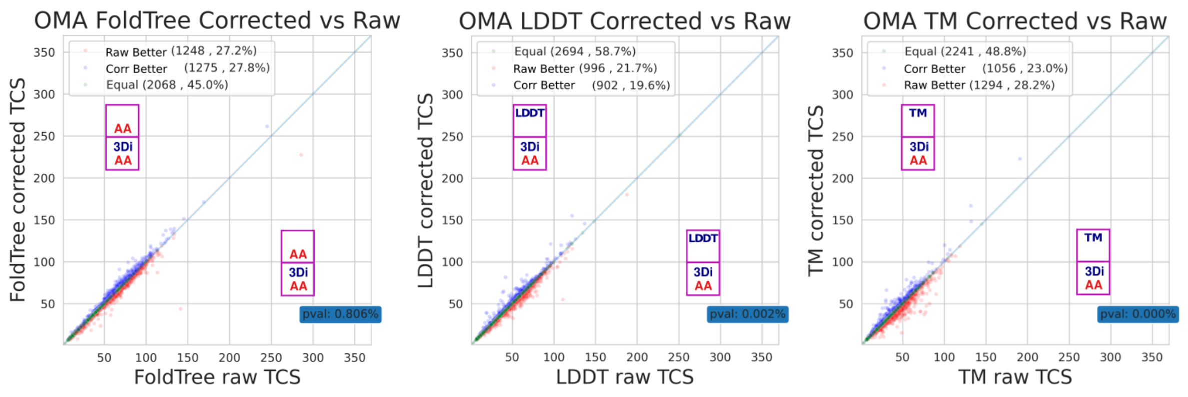


**Supplementary Figure 8.** The Effect of statistical corrections of structural distance metrics and the FoldTree score has a negligible impact on TCS differences in the OMA dataset. Each column of these pairwise plots shows the effect of correcting the distance metric on a TCS comparison between maximum likelihood trees and a structurally informed distance tree. The results presented in **Figure 1** are from the statistically corrected versions of the structurally informed distance trees.

We also compared TCS scores of OMA and CATH families using distance based structural distance metrics and the partition model shown in Puente-Leliere et al. (Puente-Lelievre et al.). The approach uses a partition of amino acid characters alongside a partition of 3di characters in order to infer tree topologies (as outlined in **Supp figure 1**). This appears to improve the TCS of the approach compared to exclusively using amino acids in a maximum likelihood framework when building trees for distantly related sequences. However, at shorter evolutionary distances, like those found in the OMA dataset, the approach does not appear to outperform the amino acid-based maximum likelihood trees.

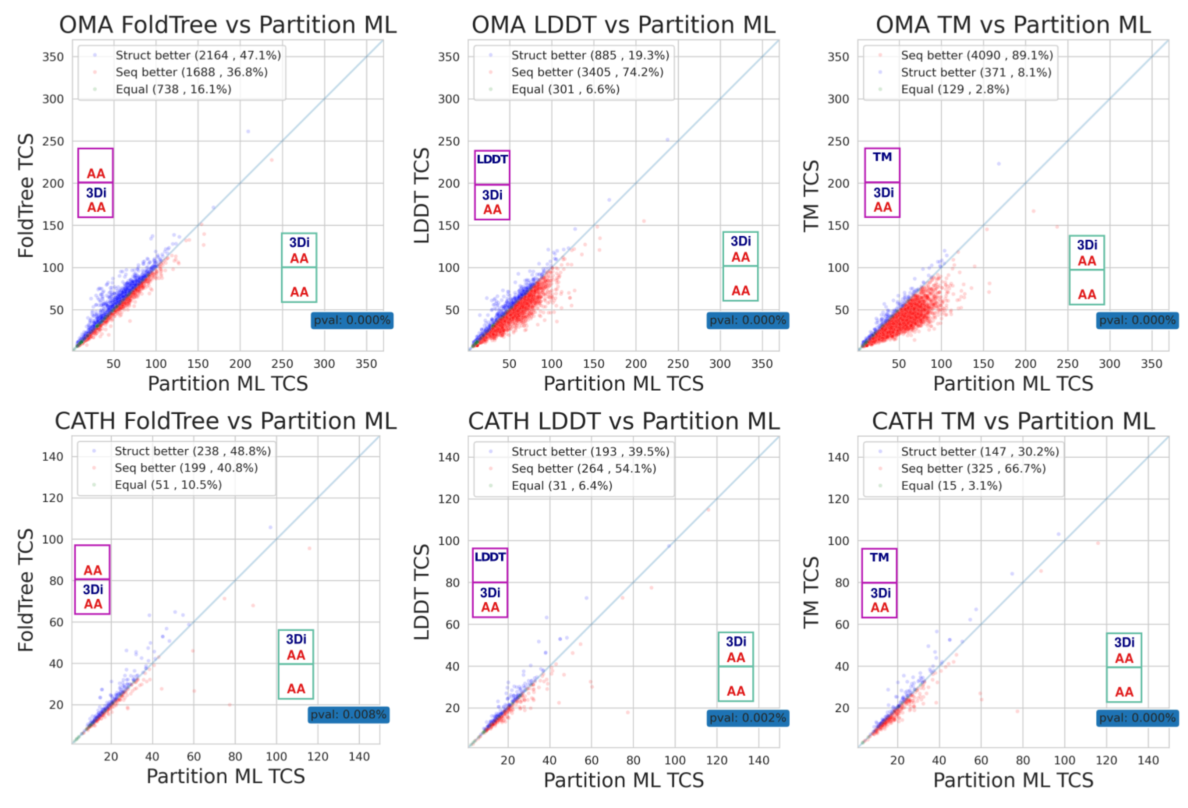


**Supplementary Figure 9. FoldTree and structural distance metric compared to the partition model approach presented in Puente-Lelievre et al.**(Puente-Lelievre et al.) FoldTree and structural distance metric-based trees are compared against a maximum likelihood model incorporating structural characters through the use of a partition model. This approach provides the evolutionary model with alignment characters which undergo a slower rate of change than amino acids. This has a positive effect on TCS values as the evolutionary divergence permitted between families increases from the OMA to the CATH dataset ( **Figure 1** ). In a pairwise comparison between the partitioned maximum likelihood approach to geometric structural distances or the FoldTree metric, we observe that, overall, FoldTree still provides the most coherent topologies.

After finding that structural quality impacted the results using our regression model (**Supplementary Figure 5**) we repeated tree building experiments after having implemented a structural quality filter which removes input proteins based on their pLDDT characteristics. The TCS comparison between sequenced-based trees and the FoldTree metric is shown in **Figure 1**. Indeed, the proportion of trees where FoldTree provides more taxonomically consistent topologies increases with the use of the filter. Below we also show the effect of filtering on FoldTree and the LDDT and TM-based based trees.

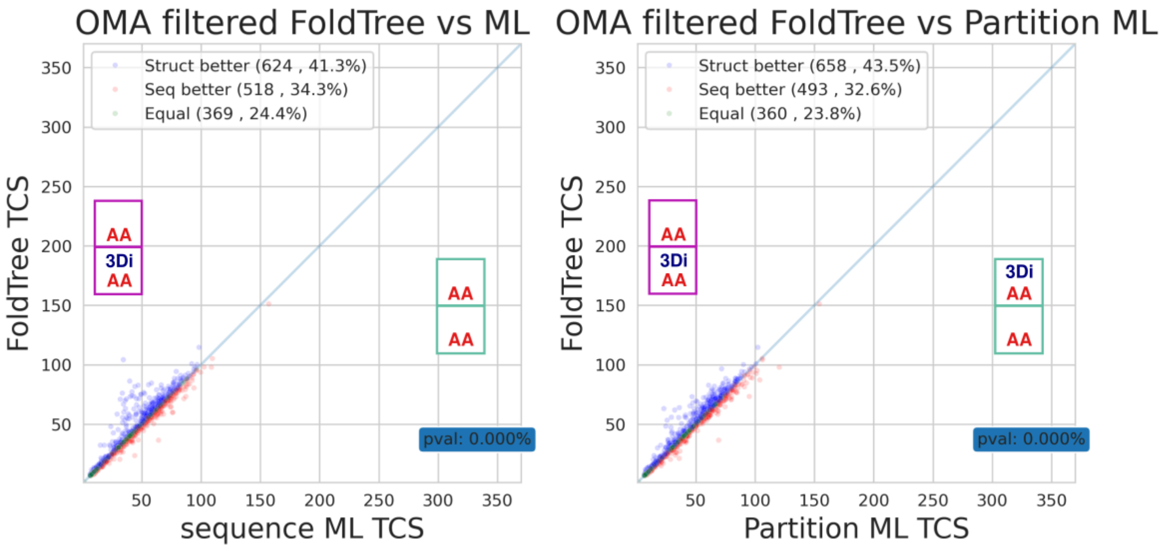


**Supplementary Figure 10. Effect of filtering input protein sets based on structural quality. a.** Performance for FoldTree and structural distance-based trees benefit from the removal of families with mean pLDDT across all proteins less than 60 and a low structural quality variance within each family ( less than 10). The proportion of trees with equal or higher TCS values, when compared to standard or structurally informed maximum likelihood trees, is greater when families with poor structure quality are removed from the dataset. Misfolded regions in alphafold structures may interfere with the inference of structural characters. This in turn may negatively affect the structurally informed pairwise alignments used as input to FoldTree. The advantage appears slightly less pronounced in the case of structurally informed maximum likelihood trees.

We also investigated whether or not the inferred time of divergence had any impact on the difference in quality of the inferred trees. We chose subsets of the OMA dataset containing families that were inferred to have emerged at specific taxonomic levels. We chose to consider HOGs that were inferred to have diverged at the Bacteria, Bilateria, Chordata, Dikarya, Eukaryota, Eumetazoa, Euteleostomi, Fungi, LUCA, Opisthokonta, and Tetrapoda levels. When considering subsets of the OMA data, it becomes apparent that some parts of the taxonomy are better resolved than others and this information is reflected in the taxonomic lineage information provided by uniprot and subsequently in the normalized TCS of trees within lineages that have a particularly poor or rich definition. However, even in recently emerged lineages such as chordata, we see that FoldTree still outperforms sequence-based methods. In many of these subsets, divergence times are low and well within the ‘wheelhouse’ of sequence-based phylogenetics. Even in this case, structure trees are, at worst, equivalent in their ability to provide topologies with taxonomic signal or, at best, slightly advantaged.

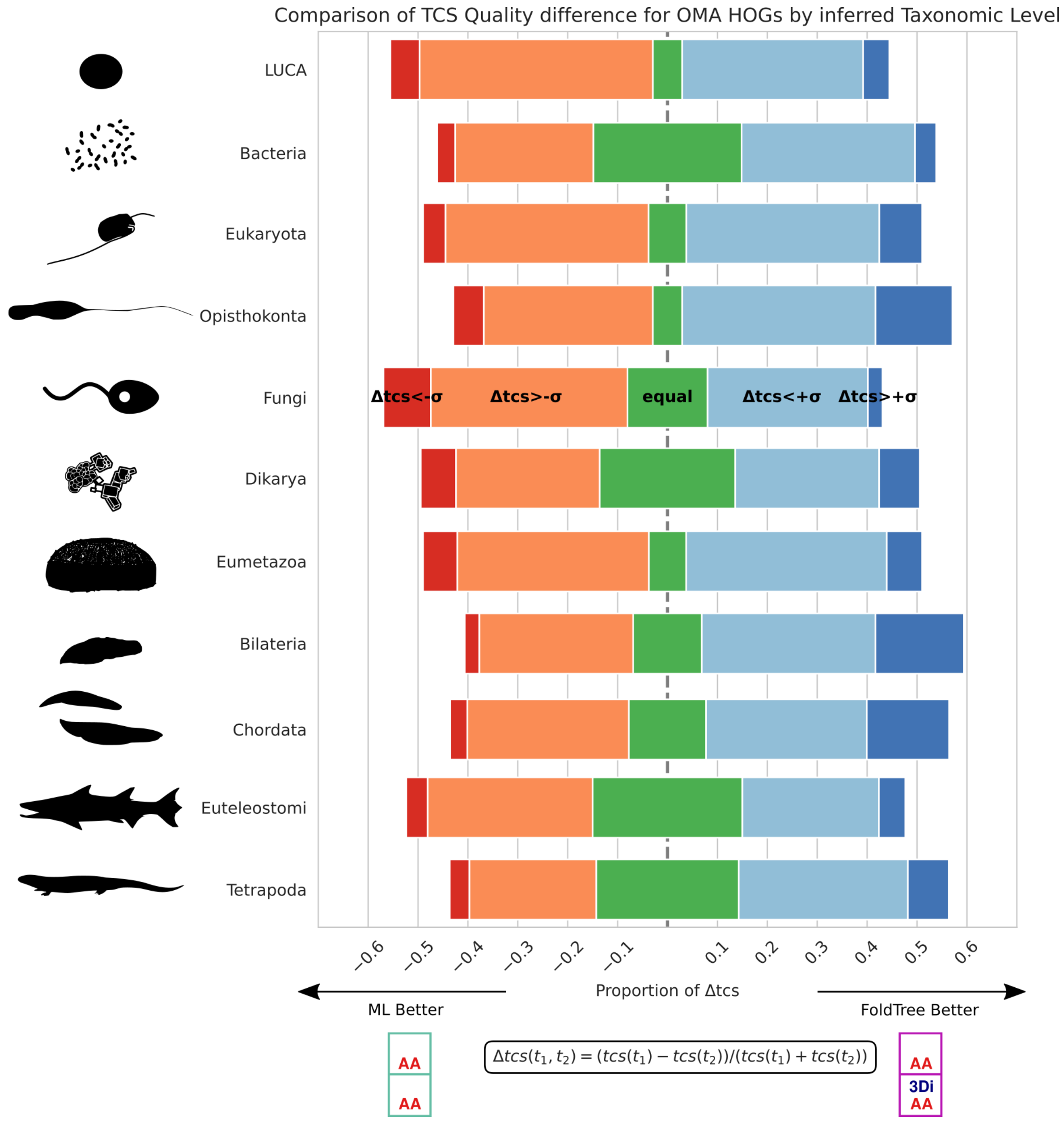


**Supplementary Figure 11.** To compare the divergence in topology quality across all families we calculated the difference in TCS scores between FoldTree and sequence-based maximum likelihood trees divided by the sum of both TCS values. Values of this score that were above one standard deviation were considered to correspond to cases where FoldTree topologies were ‘much better’ or ‘much worse’. The input dataset of OMA families was split into subsets that were inferred to have emerged at different taxonomic levels in the unfiltered OMA dataset. The inferred time of emergence is not correlated to divergence on the protein level since some protein families are under tighter evolutionary constraints than others. Interestingly, protein families that are inferred to have emerged at the LUCA level using sequence-based homology detection appear to not have diverged greatly and do not benefit from structurally informed tree building. This may be due to the fact that they are under extreme evolutionary pressure and any modifications to this set could result in non-viability.

In order to determine the effect that the observable difference in alignment quality noted in supplementary Figures 4 and 5 were having on the final topologies, we divided the OMA dataset into quartiles as a function of mean alignment pairwise distance for each family. We then compared the TCS values for all families within each quartile. The results are shown below in **Supp Figure 12**.

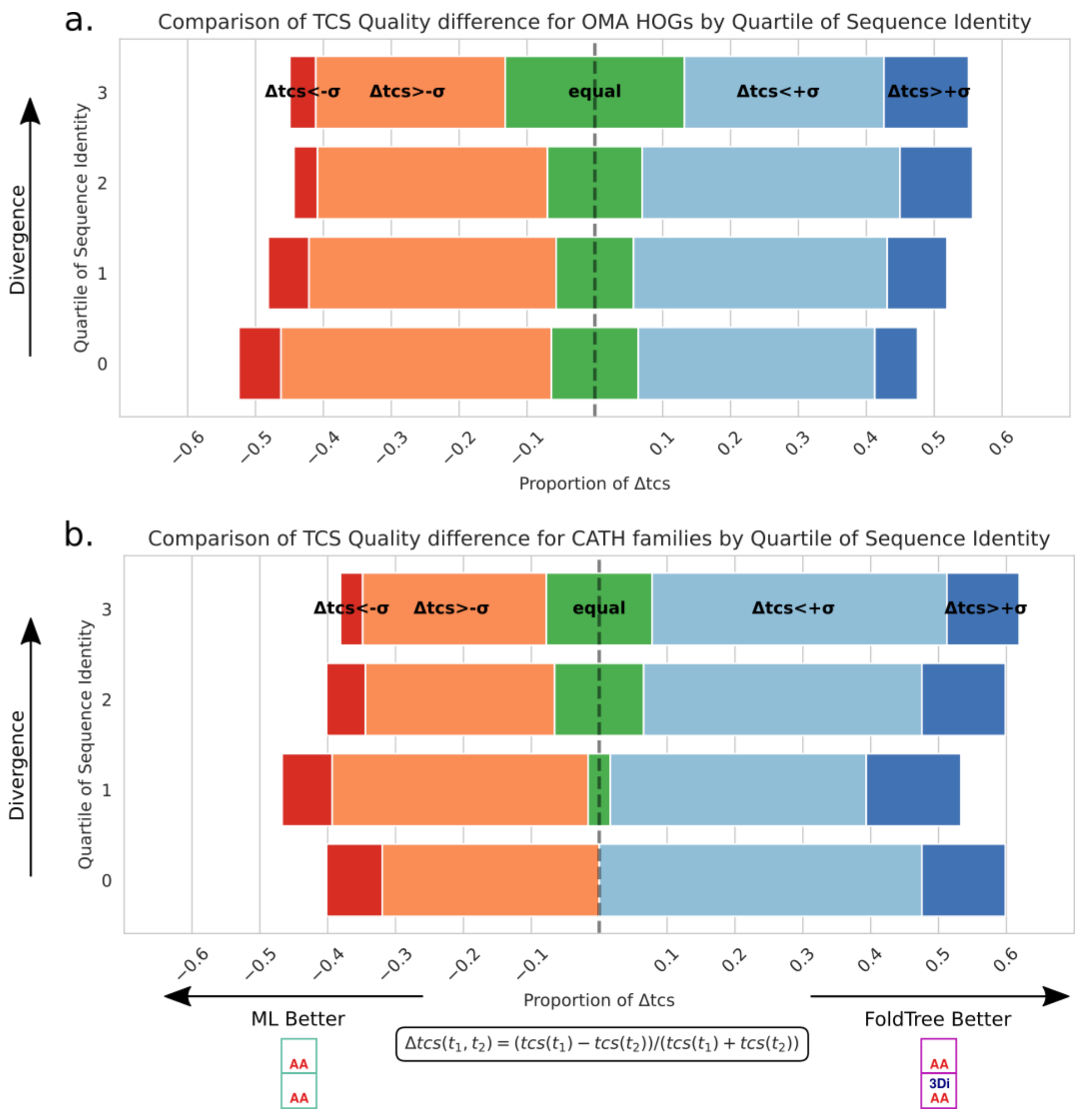


**Supplementary Figure 12**. **a )** To compare the divergence in topology quality across all families we calculated the difference in TCS scores divided by the sum of both TCS values. Differences in this score that were above one standard deviation were considered to correspond to cases where FoldTree topologies were ‘much better’ or ‘much worse’

and colored in dark red or blue. We split the TCS comparisons between FoldTree and Sequence-based trees of OMA families into quartiles as a function of their mean pairwise identity after amino acid-based alignment. As alignment becomes more difficult and mean pairwise identity drops, we see the differences in tree quality between FoldTree and Sequence-based trees become more pronounced. **b)** We repeated the analysis using CATH families and found a similar trend where more divergent amino acid alignments were associated with a higher proportion of trees where FoldTree had a 'better' or 'much better' topology.

This highlights that FoldTree's ideal use case when compared to a standard maximum likelihood sequence tree is when the input set of homologues is very divergent and alignment quality is poor.

Finally, we present our results on the CATH dataset using corrected structure-based trees. The results below in **Supp Figure 13.a** show the TCS comparison of FoldTree with the partition model method devised by Puente-Lelievre et al. as well as typical maximum likelihood trees. In **Supp Figure 13.b** we can see that FoldTree provides the lowest root to tip variance on tree topologies, followed by LDDT, Partition ML, TM and, lastly, sequence based trees. This result mirrors the outcome shown in **Figure 1.d** and **Figure 1.e** but we can see now that this topological variance is much higher in sequence-based trees while the FoldTree metric retains a similar distribution across the OMA and CATH datasets. When considering both the adherence to a molecular clock and TCS differences in the CATH dataset, we see that including structural characters in the partition model has greatly improved the clock adherence of partition model trees compared to amino acid maximum likelihood trees. This is to be expected due to the low 'mutation rate' of the structural 3di characters. However, it is unclear what meaning the branch lengths have in this context since character changes in the 3di alphabet do not have the same meaning as an amino acid character change.

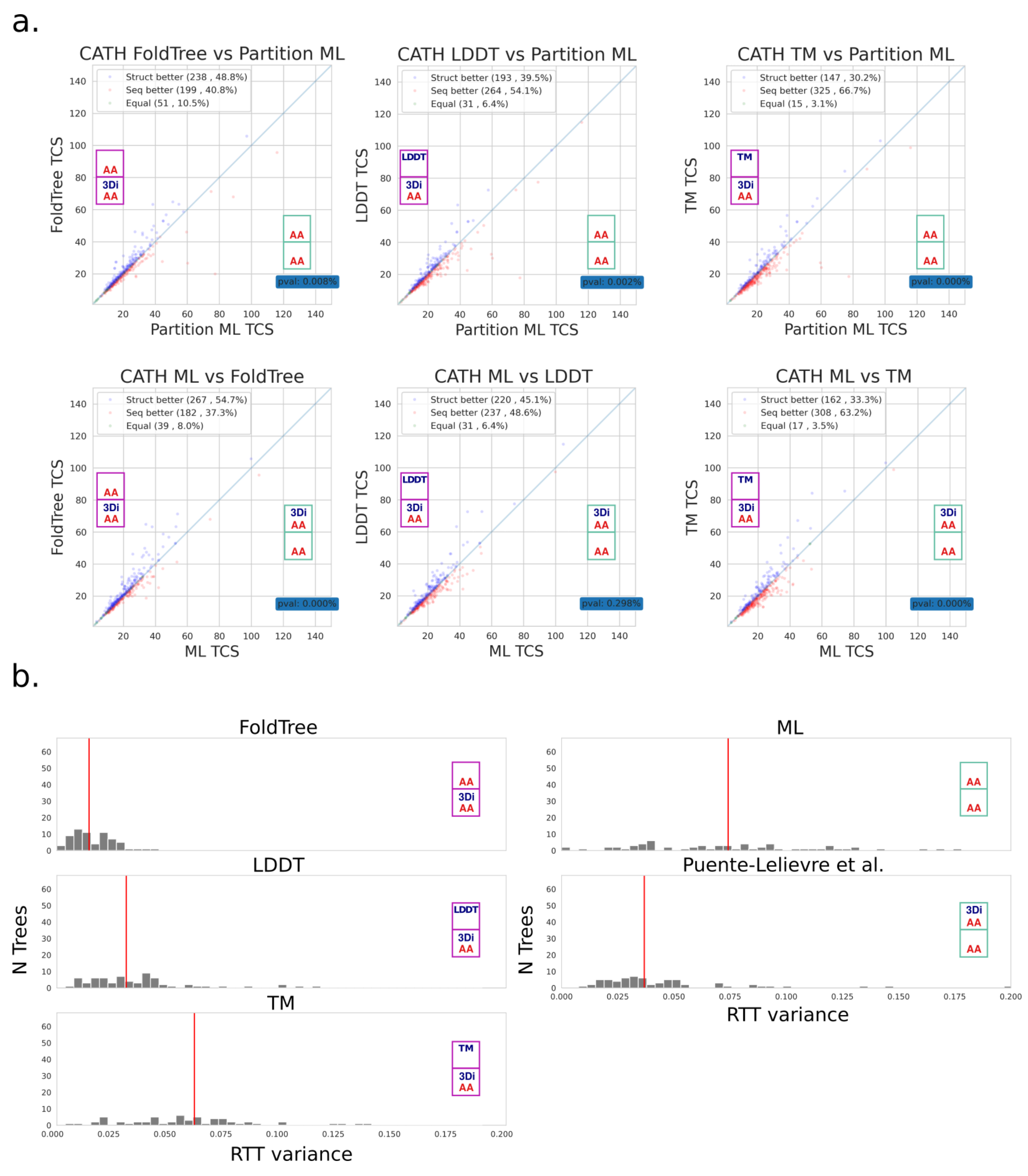


**Supplementary Figure 13. a)** TCS score comparison of structural distance metrics and Sequence-based and partition model trees against FoldTree and structural distance metrics for CATH families. **b)** Molecular clock adherence was quantified for the CATH using the same root to tip variance metric as in **Figure 1** (described in methods). The median value is shown with a red line. The statistically corrected FoldTree metric again shows lower root to tip distance variance when compared to the other methods.

When considering individual families at such large evolutionary distances, just as in our case study with RNNPA shown in **Figure 2,** it is advisable to interpret these phylogenies of very divergent homologues with a grain of salt; using orthogonal sources of information such as the taxonomic distribution of each clade, domain architecture or experimental data on the function of different representatives throughout a phylogeny

#### Astral species tree quartet support

The benchmark presented in the main text and in the supplementary figures 6-13 is designed to give more weight to correct topology deeper within the tree. To provide a more balanced perspective and give an idea of FoldTree's performance compared to maximum likelihood methods on the finer grained topology closer to the leaves of inferred trees, we used Astral to benchmark the quartet support of taxonomic trees containing all of the species within each family benchmarked. A thorough explanation of the Astral-Pro algorithm and its measure of quartet support can be found in the original manuscript (Zhang and Mirarab). In short, branches of the species tree with higher quartet support are more represented within the proposed topology for a given gene tree. We averaged the quartet support for all branches of the species trees corresponding to input protein sets for each family-wise comparison (**Supp Methods**). This measure of quartet support does not weigh the deeper branches of the tree more heavily. The Astral-Pro algorithm also infers the number of losses and duplications implied by a tree topology when compared to the species taxonomy. Below, we show these two benchmarks for the OMA dataset. FoldTree performs slightly worse overall, although it is worth noting that it does appear to outperform maximum likelihood trees in a little more than a third of topologies generated and is equivalent in a quarter. The variability observed in tree quality for individual families again highlights the need for caution in interpreting phylogenetic results and orthogonal analysis and data in determining which tree topology and implied evolutionary hypothesis is the most credible in light of the available evidence.

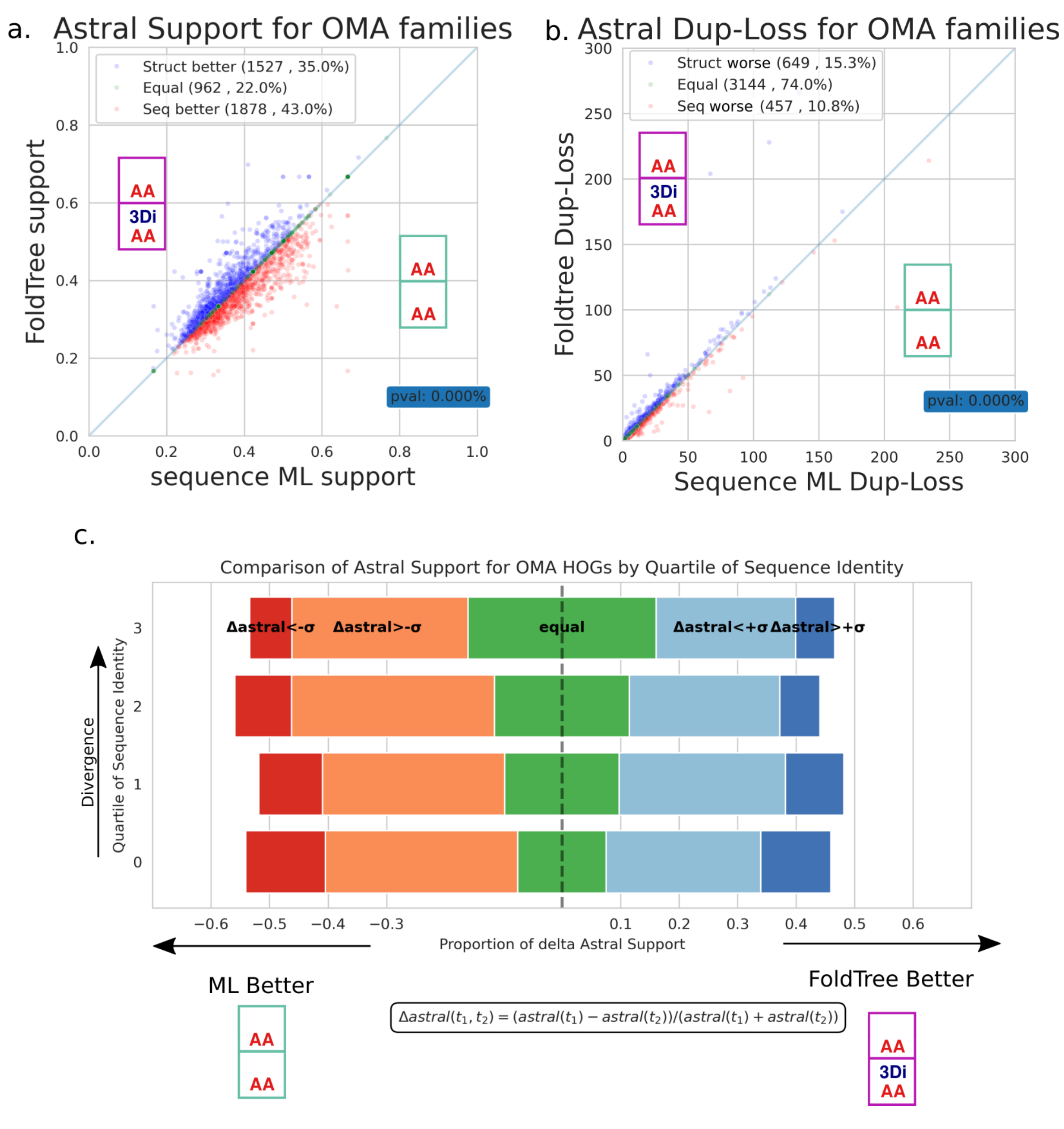


**Supplementary Figure 14.** **a)** FoldTree topologies are compared to maximum likelihood phylogenies generated using ASTRAL quartet support for species tree branches. The average support for all species tree branches is shown on the x and y axes for each family. We again observe a slight margin in favor of sequence-based trees. **b)** ASTRAL infers implied loss and duplication events in the evolutionary history of an input family by comparing a phylogeny to the species tree. Here we present the number of implied loss and duplication events for all families in the OMA dataset. A more parsimonious tree would presumably show less implied events. In this experiment only 26% of trees have a difference in terms of implied loss and duplication events with a slight margin favoring sequence-based trees. **c)** We divided the OMA dataset into quartiles of mean pairwise identity after multiple sequence alignment using amino acids. We defined the Astral comparison score as the difference of two mean support values for an input family divided by the sum of the mean support values. Any values above or below one standard deviation of this comparison score were considered ‘much worse or ‘much better’.

We repeated the Astral-Pro benchmark on the CATH dataset. Since protein families in this dataset were filtered to one representative entry per genus to avoid redundancy ( **supp methods** ), the topologies generated in this family cannot provide information relative to losses and duplications. However, the support of species tree branches can still be computed. In the comparison between maximum likelihood methods and FoldTree, at these larger evolutionary distances, the support across all quartets appears equivalent.

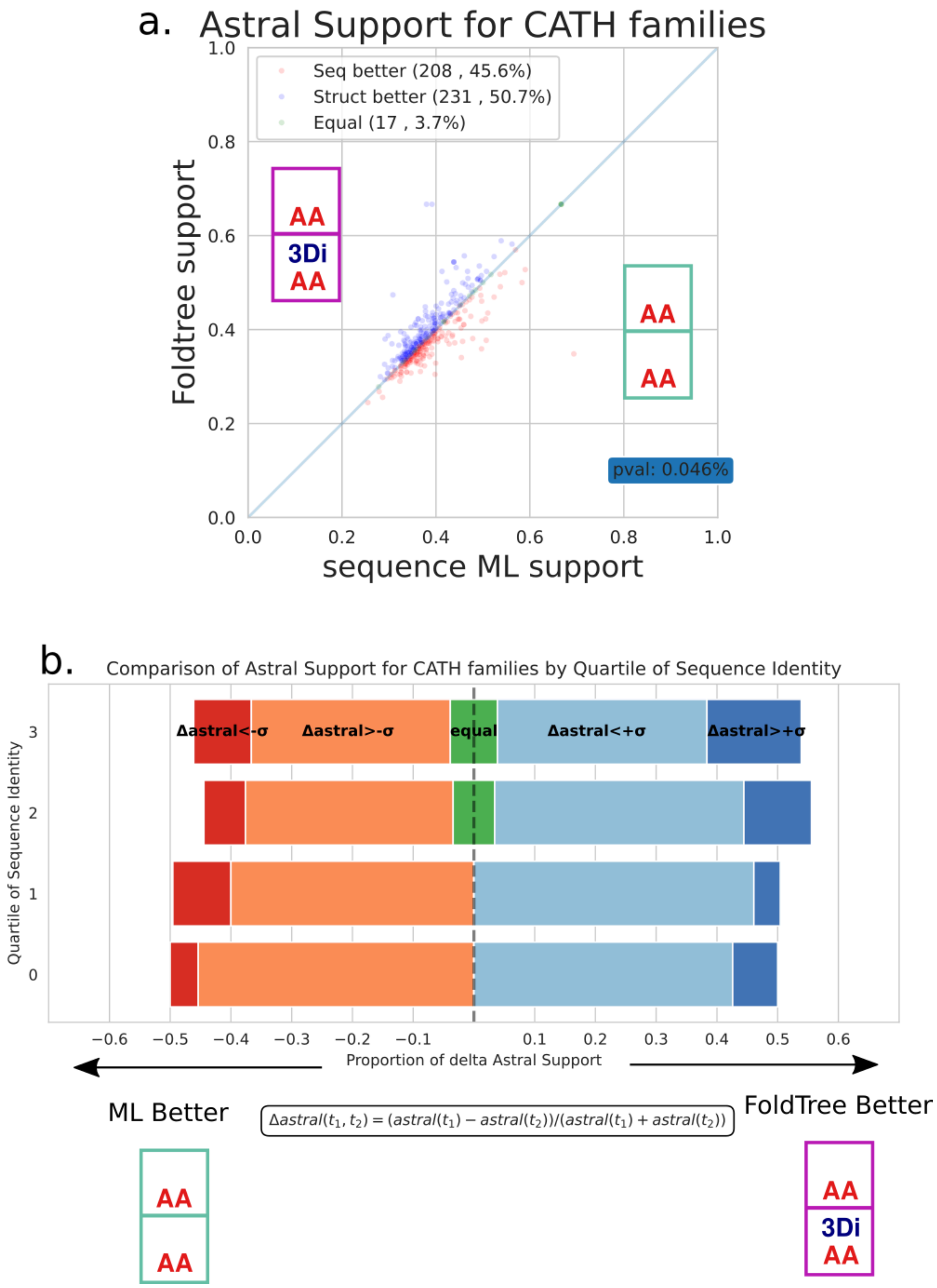


**Supplementary Figure 15. a)** We compared mean ASTRAL support for all families in the CATH dataset between sequence-based maximum likelihood trees and FoldTree. In contrast with the small margin in favor of sequence-based methods in the ASTRAL benchmark on all OMA families, we can observe the opposite in CATH families. **b)** We divided the CATH dataset into quartiles of mean pairwise identity after multiple sequence alignment using amino acids. As alignment pairwise identities decrease, we see the proportion of FoldTree trees with much better topologies increase. We defined the Astral comparison score as the difference of two mean branch support values for an input family divided by the sum of these mean support values. Any values above or below one standard deviation of this comparison score were considered ‘much worse or ‘much better’.

These observations highlight the limitations of FoldTree when dealing with the shorter evolutionary distances and relationships present closer to the leaves of the trees inferred. Maximum likelihood phylogenetics appears to have an edge when considering these shorter evolutionary distances and efforts need to be made to come up with models that can incorporate the strengths of both approaches. However, when considering families which are constructed using structural homology and permit a larger divergence of sequence identity within the family, structural phylogenetics provides topologies which are more consistent with the species tree. In these cases, both methods should be used and the final topology should be corroborated with orthogonal data, as there is no one size fits all solution. This being said, we do notice that the proportion of FoldTree trees with much better topologies, in terms of TCS and ASTRAL support when compared to the species tree, tends to increase with mean sequence divergence. This observation highlights FoldTree’s strengths and ideal use case: creating phylogenies for protein families which have diverged beyond where typical sequence-based phylogenetic methods are applicable.

Sequence-based phylogeny of RRNPPA family.

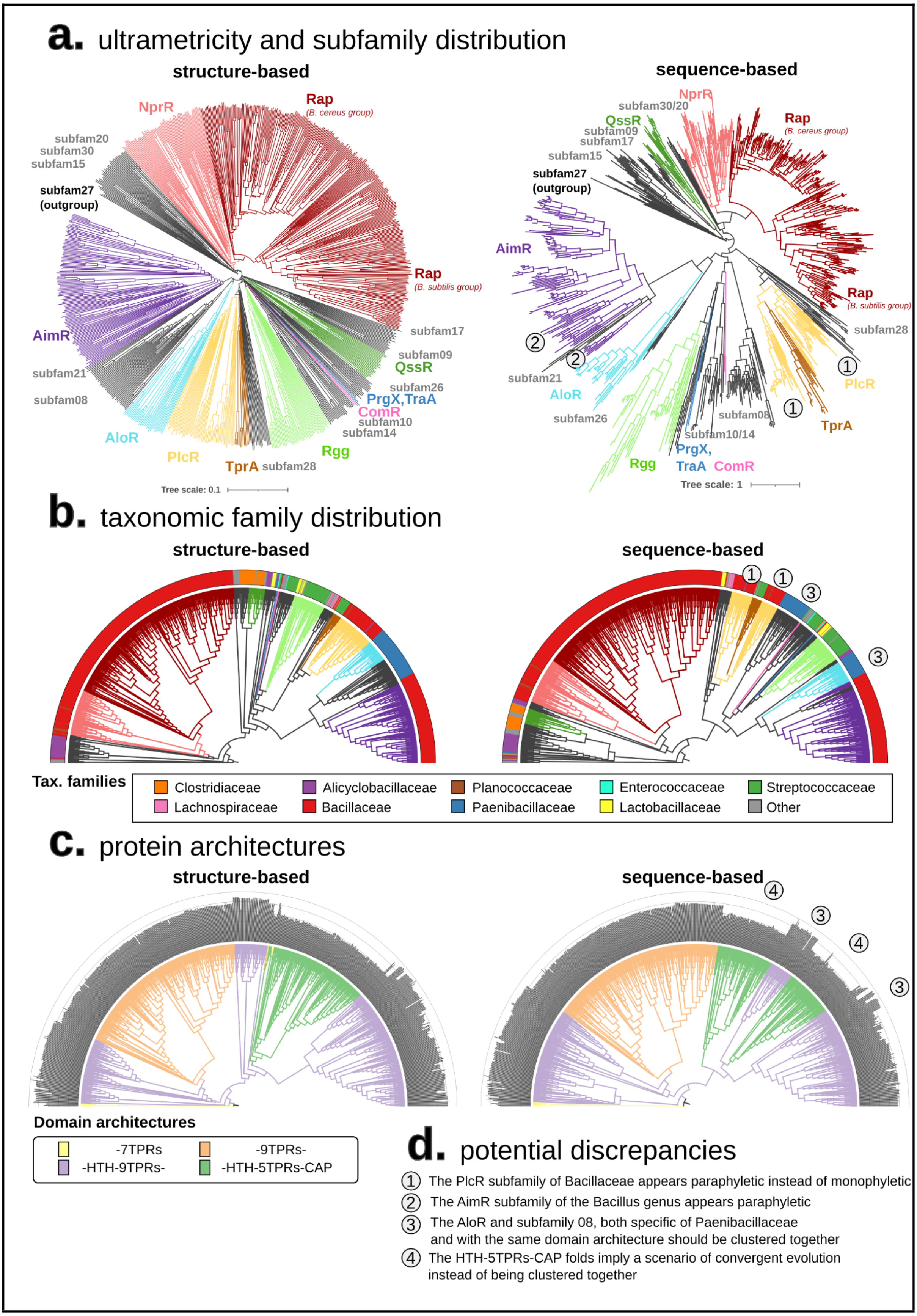

**Supplementary Figure 16.** Comparison of structure- and sequence-based phylogenies of RRNPPA family.

To appreciate the relevance of FoldTree for the reconstruction of the phylogeny of the divergent RRNPPA family, we compared its output tree with that of a state-of-the-art sequence-based pipeline consisting of an alignment with mafft (–iterate 1000 and –localpairs options for high accuracy), a trimming with TrimAl (option –automated 1 optimized for ML reconstruction), and a tree inference with IQTREE with 1000 bootstraps under the LG+G model of amino acid substitution (**Supplementary Figure 16**). For consistency, both trees were rooted in subfamily 27 of *Anoxybacter fermentans* (MAD root of the structural tree). While we do not assert this as the actual root of the phylogeny, it was chosen to facilitate tree comparisons. While the structure-based tree yields uniform root-to-tips distances, these distances appear strikingly more variable on the sequence-based tree (**Supplementary Figure 16A**). Some other discrepancies can be observed on the sequence-based tree. For instance, the AimR and PlcR subfamilies are not monophyletic. Indeed, the chromosomal systems of *Alkalihalobacilli* from subfamily 21 branches from within the clade of AimR receptors of *Bacilli* temperate phages (**Supplementary Figure 16A-B**) and the TprA subfamily of *Streptococcaceae* branches from within the PlcR clade of *Bacillaceae*. This is inconsistent with the literature (Neiditch et al.; Stokar-Avihail et al.) and not parsimonious in terms of taxonomic distribution. But most importantly, the main discrepancy of the sequence-based tree is the wrong placement of the subfamily 08 of *Paenibacillaceae* as a sister clade of Rgg-ComR-PrgX, with PlcR-TrpA as their outgroup (**Supplementary Figure 16A**). This is not parsimonious in two regards: first, it implies that HTH-5TPRs architectures emerged two independent times (**Supplementary Figure 16C**) and second, it implies a paraphyly of communication systems from *Paenibacillaceae* (**Supplementary Figure 16C**). In contrast, in the structure-based tree, subfamily 08 is a sister clade of AloR, consistent with their common architecture (HTH-9TRPS) and their common specificity to the *Paenibacillaceae* taxonomic family. This placement, as opposed to the sequence-based tree, is more parsimonious as it implies a single emergence of the HTH-9TRPs and HTH-5TPRs folds.

#### Supplementary references


Mutti, Giacomo, et al. “Newly Developed Structure-Based Methods Do Not Outperform Standard Sequence-Based Methods for Large-Scale Phylogenomics.” *Bioinformatics*, biorxiv;2024.08.02.606352v1, bioRxiv, 6 Aug. 2024, https://www.biorxiv.org/content/10.1101/2024.08.02.606352v1.full.pdf.

Neiditch, Matthew B., et al. “Genetic and Structural Analyses of RRNPP Intercellular Peptide Signaling of Gram-Positive Bacteria.” *Annual Review of Genetics*, vol. 51, Nov. 2017, pp. 311–33, https://doi.org/10.1146/annurev-genet-120116-023507.

Øksendal, Bernt. *Stochastic Differential Equations*. Springer Berlin Heidelberg, https://doi.org/10.1007/978-3-642-14394-6. Accessed 7 Mar. 2023.

Puente-Lelievre, Caroline, et al. “Tertiary-Interaction Characters Enable Fast, Model-Based Structural Phylogenetics beyond the Twilight Zone.” *bioRxiv*, 13 Dec. 2023, p. 2023.12.12.571181, https://doi.org/10.1101/2023.12.12.571181.

Rzhetsky, A., and M. Nei. “Theoretical Foundation of the Minimum-Evolution Method of Phylogenetic Inference.” *Molecular Biology and Evolution*, vol. 10, no. 5, Sept. 1993, pp. 1073–95, https://doi.org/10.1093/oxfordjournals.molbev.a040056.

Stokar-Avihail, Avigail, et al. “Widespread Utilization of Peptide Communication in Phages Infecting Soil and Pathogenic Bacteria.” *Cell Host & Microbe*, vol. 25, no. 5, May 2019, pp. 746–55.e5, https://doi.org/10.1016/j.chom.2019.03.017.

Tajima, F. “Simple Methods for Testing the Molecular Evolutionary Clock Hypothesis.” *Genetics*, vol. 135, no. 2, Oct. 1993, pp. 599–607, https://doi.org/10.1093/genetics/135.2.599.

---. “Unbiased Estimation of Evolutionary Distance between Nucleotide Sequences.” *Molecular Biology and Evolution*, vol. 10, no. 3, May 1993, pp. 677–88, https://doi.org/10.1093/oxfordjournals.molbev.a040031.

van Kempen, Michel, et al. “Fast and Accurate Protein Structure Search with Foldseek.” *Nature Biotechnology*, May 2023, https://doi.org/10.1038/s41587-023-01773-0.

Zhang, Chao, and Siavash Mirarab. “ASTRAL-Pro 2: Ultrafast Species Tree Reconstruction from Multi-Copy Gene Family Trees.” *Bioinformatics (Oxford, England)*, vol. 38, no. 21, Oct. 2022, pp. 4949–50, https://doi.org/10.1093/bioinformatics/btac620.
